# A class-specific effect of dysmyelination on the excitability of hippocampal interneurons

**DOI:** 10.1101/2023.01.10.523413

**Authors:** Delphine Pinatel, Edouard Pearlstein, Giulia Bonetto, Laurence Goutebroze, Domna Karagogeos, Valérie Crépel, Catherine Faivre-Sarrailh

## Abstract

The role of myelination for axonal conduction is well-established in projection neurons but little is known about its significance in GABAergic interneurons. Myelination is discontinuous along interneuron axons and the mechanisms controlling myelin patterning and segregation of ion channels at the nodes of Ranvier have not been elucidated. Protein 4.1B is implicated in the organization of the nodes of Ranvier as a linker between paranodal and juxtaparanodal membrane proteins to the spectrin cytoskeleton. In the present study, 4.1B KO mice are used as a genetic model to analyze the functional role of myelin in Lhx6-positive parvalbumin and somatostatin neurons, two major classes of GABAergic neurons in the hippocampus. We show that deletion of 4.1B induces disruption of juxtaparanodal K^+^ channel clustering and mislocalization of nodal or heminodal Na^+^ channels. Strikingly, 4.1B-deficiency causes loss of myelin in GABAergic axons in the hippocampus. In particular, stratum oriens O-LM cells display severe axonal dysmyelination and a reduced excitability. This reduced excitability is associated with a decrease in occurrence probability of small amplitude synaptic inhibitory events on pyramidal cells. In contrast, stratum pyramidale fast-spiking basket cells do not appear affected. The aberrant myelination of hippocampal interneurons is also correlated with impairment of spatial memory in 4.1B KO mice. In conclusion, our results indicate a class-specific effect of dysmyelination on the excitability of hippocampal interneurons associated with a functional alteration of inhibitory drive and impairment of spatial memory.

## Introduction

Recent reports have established that subtypes of GABAergic neurons could be myelinated. Indeed, a substantial fraction of myelin, both in mouse and human neocortex, belongs to inhibitory neurons by comparison with pyramidal cells (Micheva et al., 2016; Stedehouder et al., 2017). Long-range projecting hippocampal GABAergic neurons are myelinated and play a role in the synchronization among distant brain areas (Jinno et al., 2007; Melzer et al., 2012). Surprisingly, locally-projecting interneurons such as the parvalbumin (PV) cells are frequently myelinated, independently of their morphological subtypes (i.e. basket or bi-stratified) in the mouse hippocampus. A fraction of somatostatin (SST) interneurons, which include both long-range and Oriens-Lacunosum Moleculare (O-LM) or bistratified local projecting neurons has been also reported to be myelinated (Micheva et al., 2016; Stedehouder et al., 2017, 2019). Myelin optimizes action potential (AP) propagation by forming electrical insulation and restricting ion channel distribution, but its significance is still elusive in local projecting interneurons. Oligodendrocytes wrapping of GABAergic axons may be essential for providing axonal metabolic support (Krasnow and Attwell, 2016; Philips and Rothstein, 2017). Recent reports indicate that myelination of PV interneurons is modulated by neuronal activity or sensory experience and is able to shape the inhibitory circuitry (Benamer et al., 2020; Stedehouder et al., 2018; Yang et al., 2020). However, how myelination could modulate the function of the different subtypes of interneurons remains poorly evaluated.

In the neocortex and hippocampus, GABAergic interneurons present the particularity to be covered by myelin sheaths that display a patchy distribution along the axons (Benamer et al., 2020; Micheva et al., 2016; Stedehouder et al., 2017).The discontinuous myelination of inhibitory ramified axons implies the presence of heminodal structures along mature axons. The mechanisms regulating myelin coverage as well as segregation of ion channels at heminodes and nodes of Ranvier remain to be elucidated. Remarkably, hippocampal PV and SST neurons display clusters of Na^+^ channels forming prenodes along the axons before myelination (Bonetto et al., 2019; Freeman et al., 2016). These prenodes can be induced by oligodendrocyte-secreted matrix molecules including Contactin (Dubessy et al., 2019). Such Na^+^ channel clusters along pre-myelinated inhibitory axons are associated with increased conduction velocity (Freeman et al., 2015). We also showed that the axons of hippocampal PV and SST interneurons are highly enriched before myelination in Kv1 channels that may regulate firing during development. The Kv1 channels are associated with the cell adhesion molecules Contactin2/TAG-1, Caspr2, and ADAM22 and the scaffolding protein 4.1B forming complexes in the juxtaparanodal domains of the nodes of Ranvier after myelination (Bonetto et al., 2019). It is however so far unknown whether deficiency in any of the juxtaparanodal components Contactin2/TAG-1, Caspr2 or 4.1B could affect the myelination of hippocampal interneurons. Here, we focused our study on the consequences of 4.1B deficiency.

Protein 4.1B belongs to a superfamily of proteins that share a conserved FERM domain (4.1/ ezrin/radixin/moesin) and serve as membrane cytoskeleton linkers and participate in a wide variety of cellular events such as motility and cell adhesion (Denisenko-Nehrbass et al., 2003). In myelinated axons, protein 4.1B is implicated in the organization of the nodes of Ranvier as a linker between paranodal and juxtaparanodal membrane proteins to the actin/spectrin cytoskeleton. At juxtaparanodes, 4.1B is linked to the Caspr2/Contactin2 cell adhesion complex and is required for the clustering of the Kv1 channels as reported in the PNS and CNS (Buttermore et al., 2011; Cifuentes-Diaz et al., 2011; Einheber et al., 2013; Horresh et al., 2010). At paranodes, 4.1B mediates the linkage between the Caspr/Contactin adhesion complex and ßII-spectrin and participates to the boundary with the nodal complex enriched in Nav channels and Neurofascin186 anchored to ßIV-spectrin/AnkyrinG scaffold (Brivio et al., 2017; Zonta et al., 2008). In addition to its role in the organization of ion channel domains at the nodes of Ranvier, 4.1B linked with Caspr at paranodes has been also reported to promote internodal elongation during development (Brivio et al., 2017).

We discovered that *4.1B*^-/-^ mice display selective aberrant myelination of GABAergic interneurons in the hippocampus. An intriguing question is whether hippocampal interneurons require paranodal cytoskeleton as an instructive cue regulating the extent of myelin sheath coverage. We study the cellular features of SST and PV cells in the hippocampus by examining ion channel distribution, axon initial segment (AIS) topography and electrophysiological intrinsic properties. We show that 4.1B-deficiency induces a dramatic and selective loss of myelin sheath along SST and PV axons in the stratum radiatum of CA1 hippocampus. Notably, the excitability of O-LM cells located in the stratum oriens is specifically reduced in contrast to PV basket cells not affected in the stratum pyramidale. This study reveals the functional role of myelin in affecting the excitability of subtypes of hippocampal interneurons and modulating spatial-related behavior.

## Results

### Deletion of the scaffolding protein 4.1B causes severe loss of myelination in inhibitory axons of the stratum radiatum in the hippocampus

We used the transgenic *Lhx6-tdTomato* reporter line (Jackson laboratory) to fluorescently label subtypes of GABAergic neurons. Lhx6 is a transcription factor expressed in inhibitory neuron subclasses originating from the medial ganglionic eminence, which includes all hippocampal PV and SST neurons (Liodis et al., 2007). Double-immunostaining for MBP as a myelin marker, and Caspr as a paranodal marker was performed and we evaluated that 75 ± 5 % of the myelinated axons were Lhx6-positive in the stratum radiatum at P35 (Fig. S1A, D).

We showed previously that cell adhesion and scaffolding molecules associated in complex with juxtaparanodal Kv1 are highly expressed in hippocampal GABAergic interneurons (Bonetto et al., 2019). Contactin2 is selectively expressed in SST cells in the stratum oriens and PV cells in the stratum pyramidale of the rat hippocampus (Bonetto et al., 2019). Here we ask whether deficiency in any of the juxtaparanodal components Contactin2/TAG-1, Caspr2 or 4.1B could induce dysmyelination of hippocampal interneurons. Strikingly, we observed that 4.1B KO mice displayed a severe loss of myelin sheaths in the hippocampus as shown in Fig.1. The myelin pattern was selectively altered in the hippocampus of the 4.1B KO mice, markedly in the stratum radiatum both at P35 and P70 (Fig. 1B, F and S1H) as an indication that myelination of inhibitory interneurons may be severely affected. An apparent slight dysmyelination was also observed in the molecular layer of the dentate gyrus (Fig. 1B inset and Fig. 3D). We did not observe any massive loss of myelin outside the hippocampus, as shown in myelinated tracts like the corpus callosum (Fig. 1B) or in the layers of the somato-sensory cortex (Fig. S2). In contrast, Contactin2-deficiency did not markedly impair myelination in the CA1 hippocampus as illustrated at P70 (Fig. 1D). In *Contactin2*^-/-^ mice crossed with the *Lhx6-tdTomato* reporter line, the total length of myelinated axons and the percentage of Lhx6-positive myelinated axons in the stratum radiatum were similar to control mice as quantified at P35 (Fig. S1A-D). As reported in several studies, Caspr2 KO mice show a reduced density of PV-positive interneurons in the hippocampus associated with decreased inhibitory synaptic transmission in CA1 pyramidal neurons (Paterno et al., 2021; Penagarikano et al., 2011). However, we did not observe any major alteration of myelinated PV inhibitory axons in the hippocampus at P70 (Fig. 1E and S1E, F).

**Figure 1:**
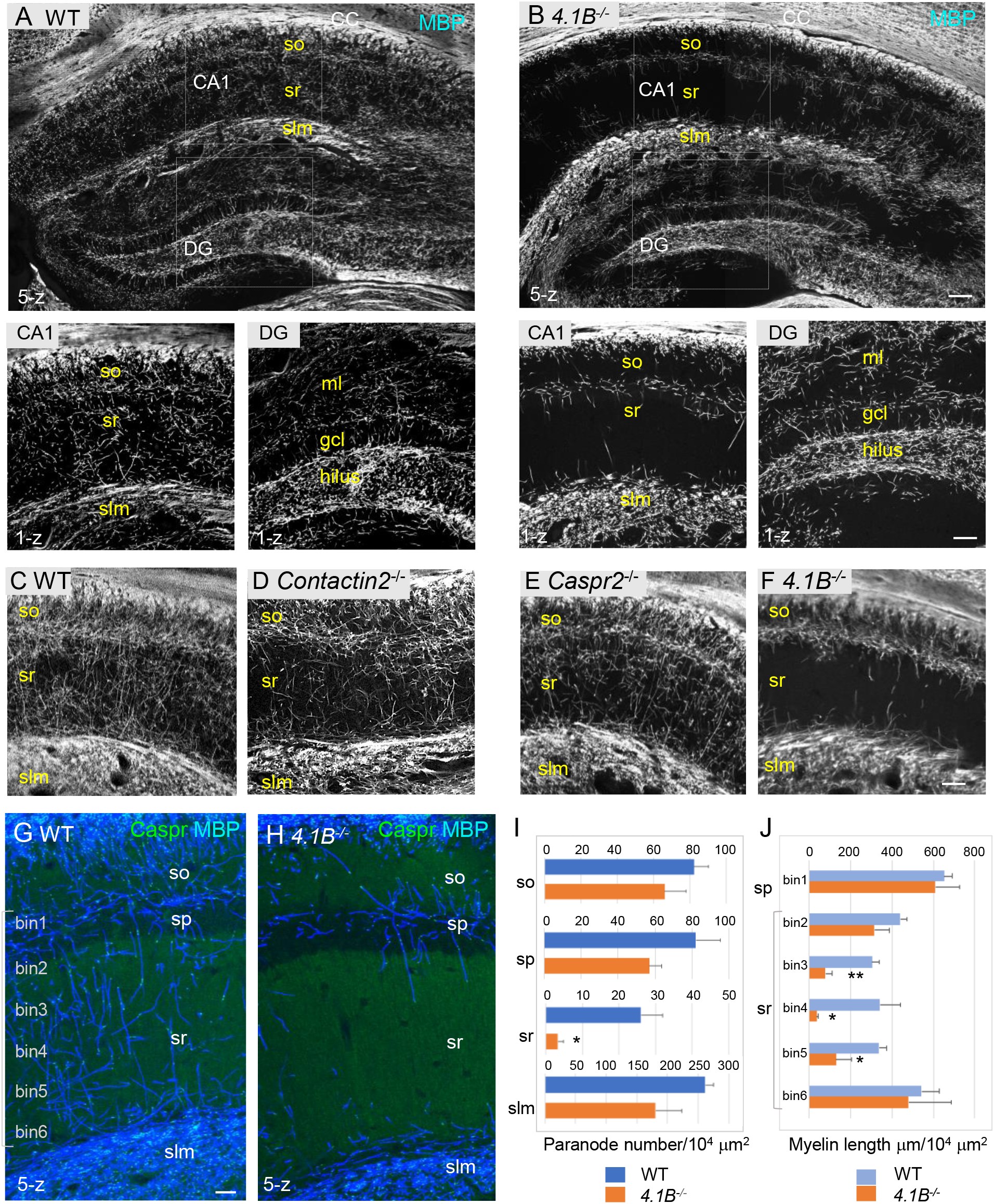
4.1B KO mice show strong alteration of myelin in the adult hippocampus. A-B: Hippocampal vibratome sections from P35 wild-type and *4.1B*^-/-^ mice immunostained for MBP as a myelin marker. Note the lack of myelin sheaths in the CA1 stratum radiatum of the *4.1B*^-/-^ hippocampus whereas myelin of the corpus callosum (CC) is preserved. Insets show high magnification of CA1 regions and dentate gyrus (DG) C-F: Hippocampus from P70 wild-type, *Contactin2*^-/-^, *Caspr2*^-/-^, and *4.1B*^-/-^ mice immunostained for MBP. Only *4.1B*^-/-^ mice show a selective and massive loss of myelin in the stratum radiatum. G, H: Double-staining for MBP (blue) and Caspr (green) as a paranodal marker. Maximum intensity of confocal images (5-z steps of 2 μm). F: Quantification of the number of paranodes/10^4^μm^2^ in the different layers of the CA1 hippocampus. G: Quantification of the total myelin length/10^4^μm^2^ in the sp (binl) and sr divided in 5 bins (40 x300 μm). Mean ± SEM of 3-4 mice for each genotype. Significant difference by comparison with wild-type: * P<0.05 and ** P<0.01 using the Student’s t-test. so: stratum oriens, sp: stratum pyramidale, sr: stratum radiatum, slm: stratum lacunosum moleculare, ml: molecular layer, gcl: granule cell layer. Bar: 200 μm in A, B; 50 μm in insets and C-F; 25 μm in G, H.

**Figure 2:**
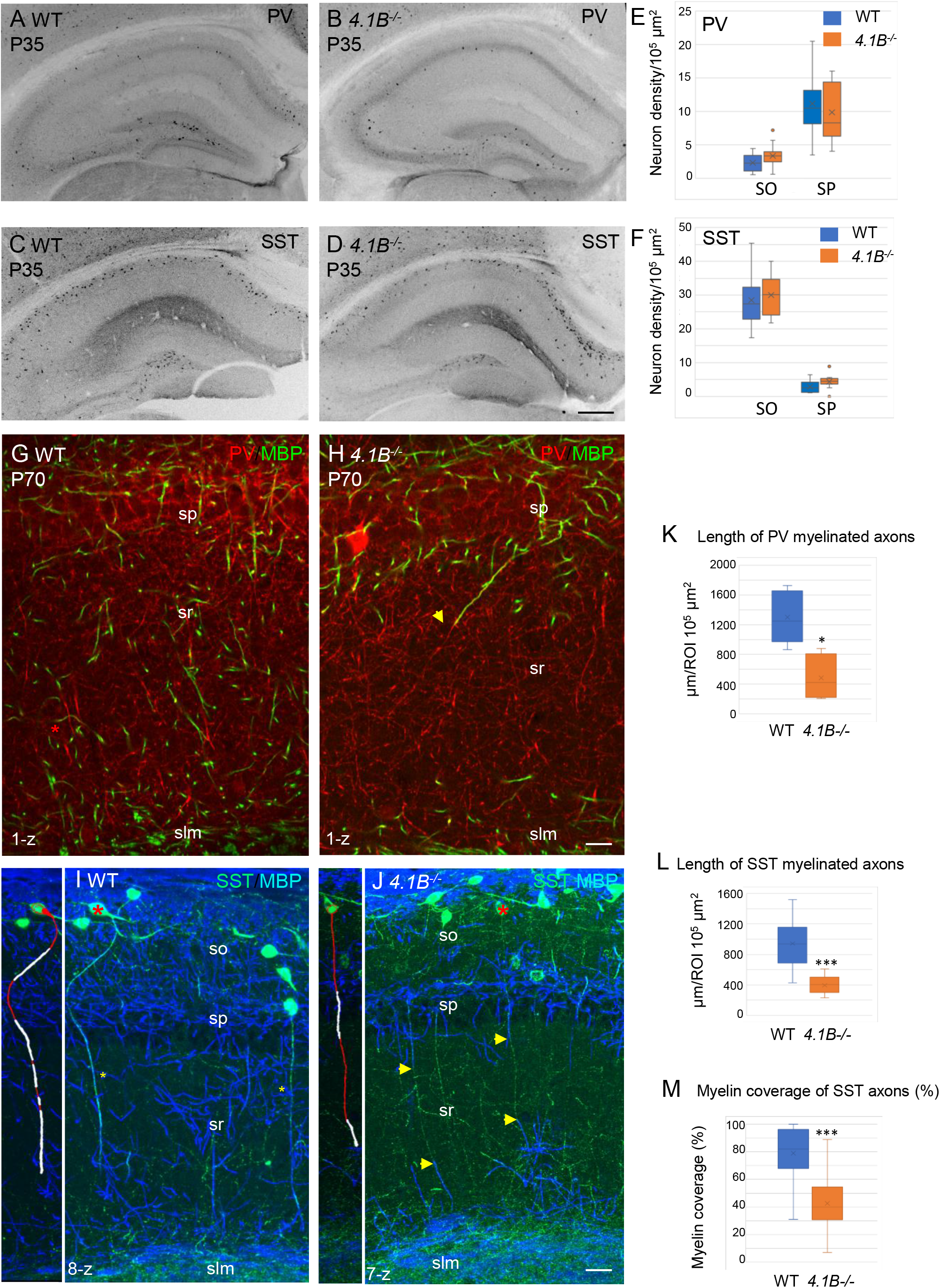
Parvalbumin and somatostatin axons are dysmyelinated in the *4.1B*^-/-^hippocampus. A-D: Hippocampal sections of wild-type and *4.1B*^-/-^ mice immunostained for PV (A, B) or SST (C, D) at P35. E, F: Quantification of the cell density/10^5^μm^2^ in the CA1 stratum oriens (SO) and stratum pyramidale (SP). No statistical difference in cell density between the genotypes (PV cell density: SO, P=0.108; SP, P=0.412 and SST cell density: SO, P=0.413; SP, P=0.057; Student’s t test; n=10-12 ROIs in hippocampal sections at 2 levels in the antero-posterior axis, 3 mice/genotype). G, H: Double-staining for PV (red) and MBP (green) at P70 showing that PV axons are present in the stratum radiatum and poorly myelinated in the *4.1B*^-/-^ mice (H). The arrow in H points to a PV axon, which is partly myelinated. K: Quantification of the total length of PV myelinated axon/10^5^μm^2^ in the stratum radiatum at P70 (n=5 ROIs, 3 mice/genotype). I, J: Double-staining for SST (green) and MBP (blue) at P35 showing O-LM SST cells in the stratum oriens extending their axon through the stratum radiatum to project into the stratum lacunosum moleculare. Insets show 3D reconstructions of myelinated O-LM neurons indicated with red asterisks (myelin in white). Note that SST axons in the stratum radiatum are fully myelinated in the WT (asterisks in I) and partly myelinated in *4.1B*^-/-^ mice (arrows in J). L, M Quantitative analyses of the total length of SST myelinated axon/10^5^μm^2^ in the stratum radiatum at P35 (n=10 ROIs, 3 mice/genotype) and the percentage of myelin coverage of individual SST axons (n=29-35 axons, 3 mice/genotype; the mean length of selected axons is 174±9 μm in WT and 197±12 μm in *4.1B*^-/-^ mice). Significant difference by comparison with wild-type: * P<0.05; *** P<0.001; Mann-Whitney test.

**Figure 3:**
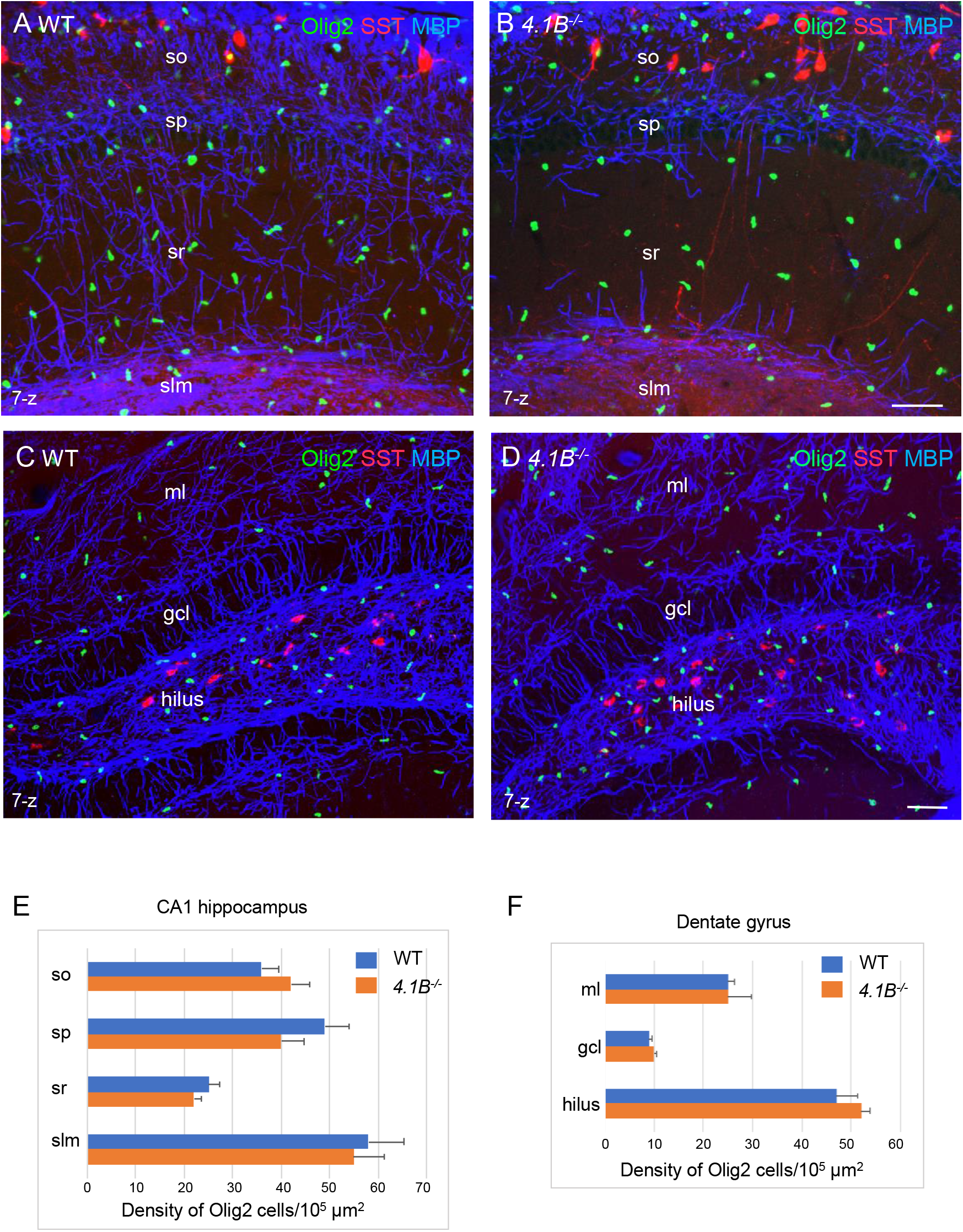
The density of oligodendroglial cells is preserved in the *4.1B*^-/-^ hippocampus. Hippocampal sections of wild-type and *4.1B*^-/-^ mice at P35 immunostained for MBP (blue), SST (red) and Olig2 (green) as a marker of the oligodendrocyte lineage. CA1 region (A, B) and dentate gyrus (C, D), maximum intensity of confocal images from 7-z steps of 2 μm. Bar: 50 μm. E, F: Quantitative analysis of the density in Olig2-positive cells/10^5^μm^2^ in the different layers of the CA1 hippocampus (n=11-12 ROIs, 3 mice/genotype) and dentate gyrus (ml: molecular layer; gcl: granular cell layer; n=3 ROIs; 3 mice/genotype). No significant difference between the genotypes (so: P=0.320; sp: P=0.090; sr: P=0.192; slm: P=0.829; ml: P=1; gcl: P=0.49; hilus: P=0.26; Student’s t test).

In the hippocampus of *4.1B*^-/-^ mice, the severe reduction of myelin sheaths occurred already at P25 in the stratum radiatum indicating that it originated from a developmental defect, taking place at an early stage of hippocampal myelination (Fig. S3). Since the extent of myelination was difficult to be quantified in regions with a dense and intermingled network of myelinated axons, the number of internodes was estimated from the number of paranodes immunolabeled for Caspr in the different layers of the CA1 hippocampus at P70 (Fig. 1G-I). The density of paranodes was significantly reduced by 88% (P=0.0147, Student’s t test) in the stratum radiatum of 4.1B-deficient mice (n=3-4 mice/genotype). The density of paranodes tended also to decrease in the stratum oriens (−18%; P=0.370), stratum pyramidale (−30%; P=0.140) and stratum lacunosum-moleculare (−31%; P=0.051). The total length of MBP-positive axonal segments of the stratum radiatum was measured at different levels starting from the pyramidal layer towards the stratum lacunosum-moleculare in the CA1 hippocampus (Fig. 1J). Myelination was significantly decreased by 75% (P=0.0015, Student’s t test) in the stratum radiatum of 4.1B-deficient mice, with a 94 % reduction in the middle region of the layer (P=0.0015). Therefore, the fact that myelin mostly belongs to Lhx6-positive axons in the stratum radiatum, suggests that myelination of inhibitory interneurons may be selectively altered in the hippocampus of 4.1B-deficient mice.

It was first critical to evaluate whether the lack of myelin could be due to a loss of GABAergic interneurons extending their axons throughout the stratum radiatum. Distinct interneurons subdivide their connections to sub-compartments of pyramidal neurons with the PV basket cells connecting the soma, the bistratified cells connecting basal and apical dendrites in the stratum oriens and radiatum, and the SST O-LM cells projecting to the distal apical dendrites in the stratum lacunosum-moleculare (Somogyi and Klausberger, 2005) (see schematic illustration Fig. 7A, H). The 4.1B KO mice did not show a reduced density of PV cells in the stratum pyramidale (Fig. 2A, B) as quantified at P35 in the CA1 region (Fig. 2E). A high density of PV axons was present in the stratum radiatum of the 4.1B-deficient mice (Fig. 2H) as observed in the wild-type (Fig. 2G), albeit they were only partly myelinated in the region bordering the stratum pyramidale (arrow in Fig. 2H) and the total length of myelinated PV axons was significantly reduced (−63%, P=0.0317, Mann-Whitney test) (Fig. 2K). The density of SST interneurons in the stratum oriens was also preserved (Fig. 2C, D, F). The axons of O-LM interneurons crossing the stratum radiatum and projecting to the stratum lacunosum-moleculare were observed in the 4.1B-deficient mice (Fig. 2J) as in the wild-type (Fig. 2I). However, O-LM axons were only partly myelinated in the lower and upper regions of the stratum radiatum (arrows in Fig. 2J) in the 4.1B KO mice with a myelin coverage of 42.3 ± 4.0% (Fig. 2M) while myelin coverage was 79.7 ± 3.4% for wild-type mice (Fig. 2I and M). The total length of myelinated SST axons was significantly reduced (−58%, P<0.0001, Mann-Whitney test) in the stratum radiatum (Fig. 2L). Thus, 4.1B-deficiency strongly reduces myelin ensheathing of PV and SST axons in the stratum radiatum.

**Figure 4:**
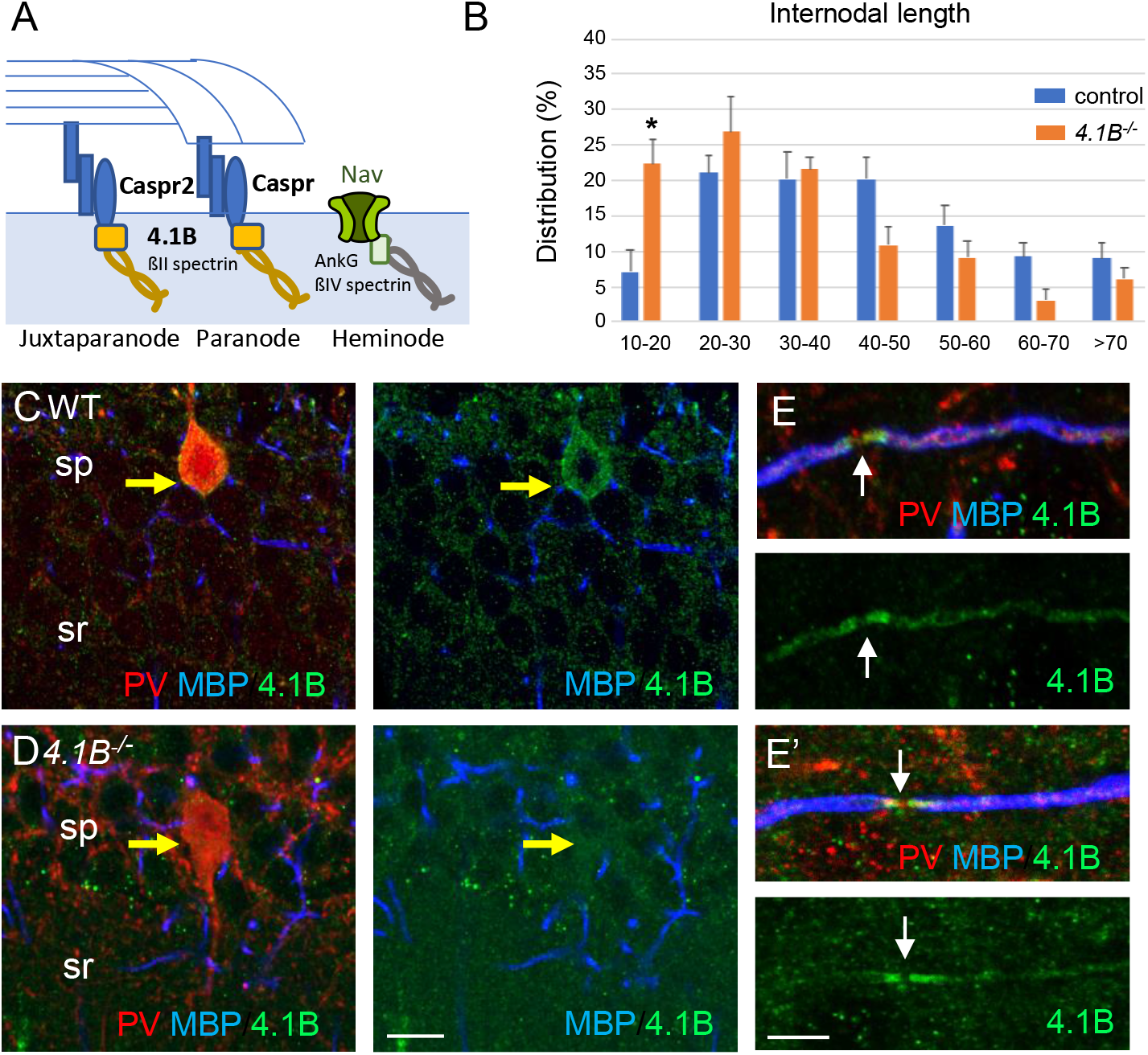
4.1B is enriched at paranodes of GABAergic axons and involved in internodal elongation. A: Schematic drawing to depict the role of 4.1B at paranodes mediating the anchorage between Caspr and the ßll-spectrin at the boundary with the nodal ßlV-spectrin. B: Distribution of internodal lengths as percentage. Quantitative analysis was performed on hippocampal sections of *Lhx6-Cre;tdTomato* control and *4.1B*^-/-^ mice at P70 immunostained for MBP and Caspr. Confocal imaging of 25 to 50-μm width z-stack were used to measure the length of myelin sheaths between two paranodes in the CA1 region (n= 275 in control and n=191 in *4.1B*^-/-^ mice). Significant difference by comparison with control: * P<0.05; Mann-Whitney test (Means ± SEM of 4 ROIs from 2 mice/genotype). C-E: Hippocampal sections of wild-type and *4.1B*^-/-^ mice at P35 immunostained for MBP (blue), PV (red) and 4.1B (green). Note that 4.1B is detected at the soma (C) and along the axons of PV interneurons, being enriched at paranodes and not detected at the nodal gap (E, E’). As a control for specificity, no staining is detected in *4.1B*^-/-^ hippocampus (D). Bar: 10 μm in C, D; 5 μm in E, E’.

**Figure 5:**
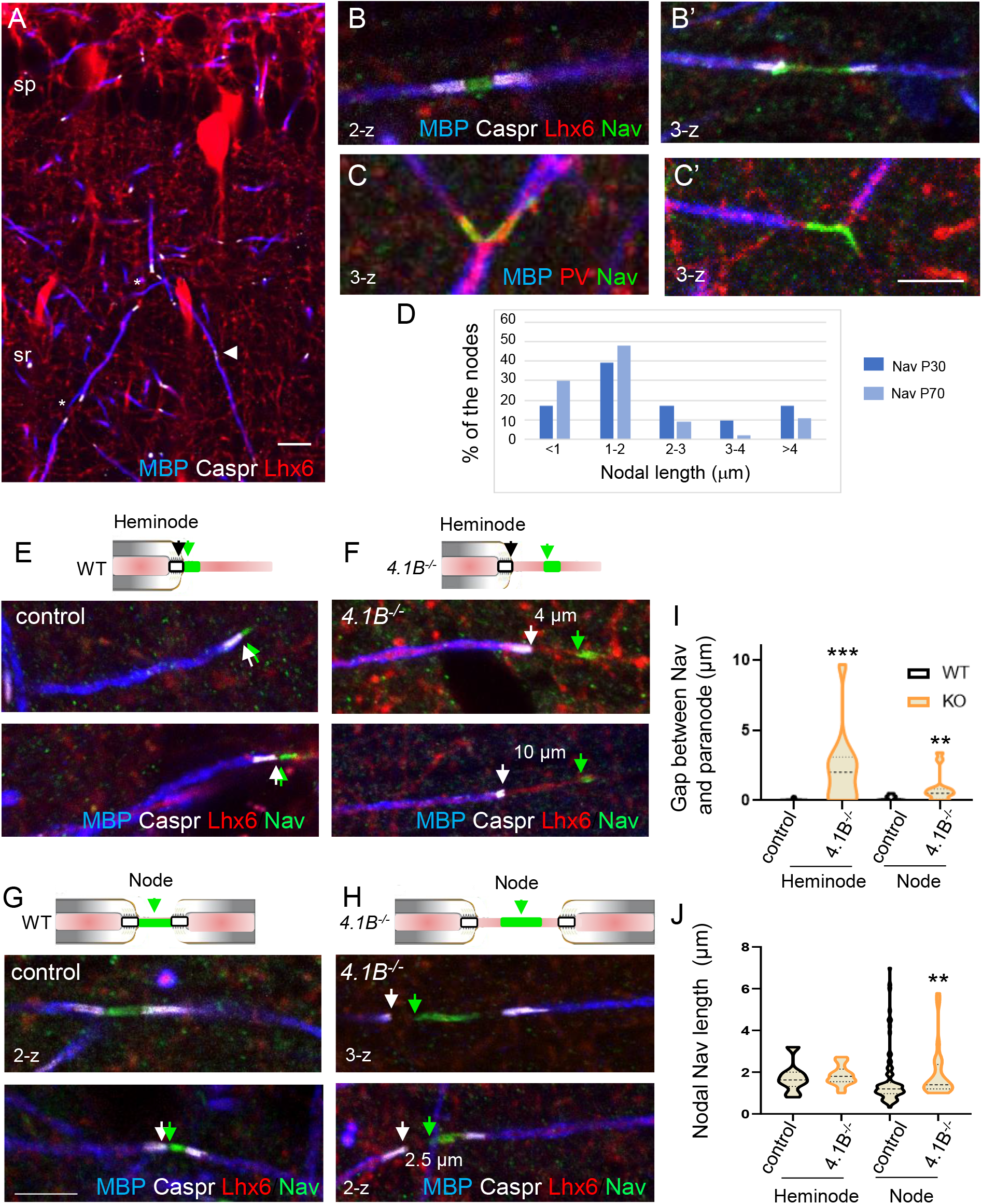
The positioning of heminodal and nodal of Nav channels is altered in 4.1B-deficient GABAergic axons. Hippocampal sections of P70 *Lhx6-Cre;tdTomato* control and *4.1B*^-/-^ mice were immunostained for MBP (blue), Caspr (white) and panNav (green). tdTomato expressed in Lhx6-positive cells or PV immunostaining (C) is in red. A: shows a branched myelinated Lhx6 axon with contiguous (arrowhead) and spaced (asterisks) internodes. Immunostaining for panNav at nodes with different lengths (B, B’) and at branch points (C, C’). D: Distribution of nodal lengths in Lhx6-positive axons at P30 and P70 in CA1 expressed as percentages. The length of nodal Nav was measured between two paranodes (n=41 at P30 and n=64 at P70). E-J: Clustering of Nav channels at the heminodes (E, F) and nodes (G, H) of control and *4.1B*^-/-^ mice in the CA1 hippocampus. Nav clusters are close to paranodes both at heminodes (E) and nodes (G) in control GABAergic axons (arrows). In contrast, a gap is frequently observed between Nav clusters and paranodes at heminodes (F) and nodes (H) in 4.1B-deficient GABAergic axons (arrows). Confocal images with maximum intensity of 2-z or 3-z steps of 1 μm. Bar: 10 μm in A; 5 μm in B, C, E-H. I: Quantitative analysis of the distance between Nav cluster and paranode at heminodes and nodes. J: Quantitative analysis of the length of nodal Nav cluster at heminodes and nodes. Significant difference by comparison with wild-type: ** P<0.01; *** P<0.001; Mann-Whitney test (heminodes: n= 25/genotype; nodes: n=87 in control and n=26 in 4.1B KO mice).

**Figure 6:**
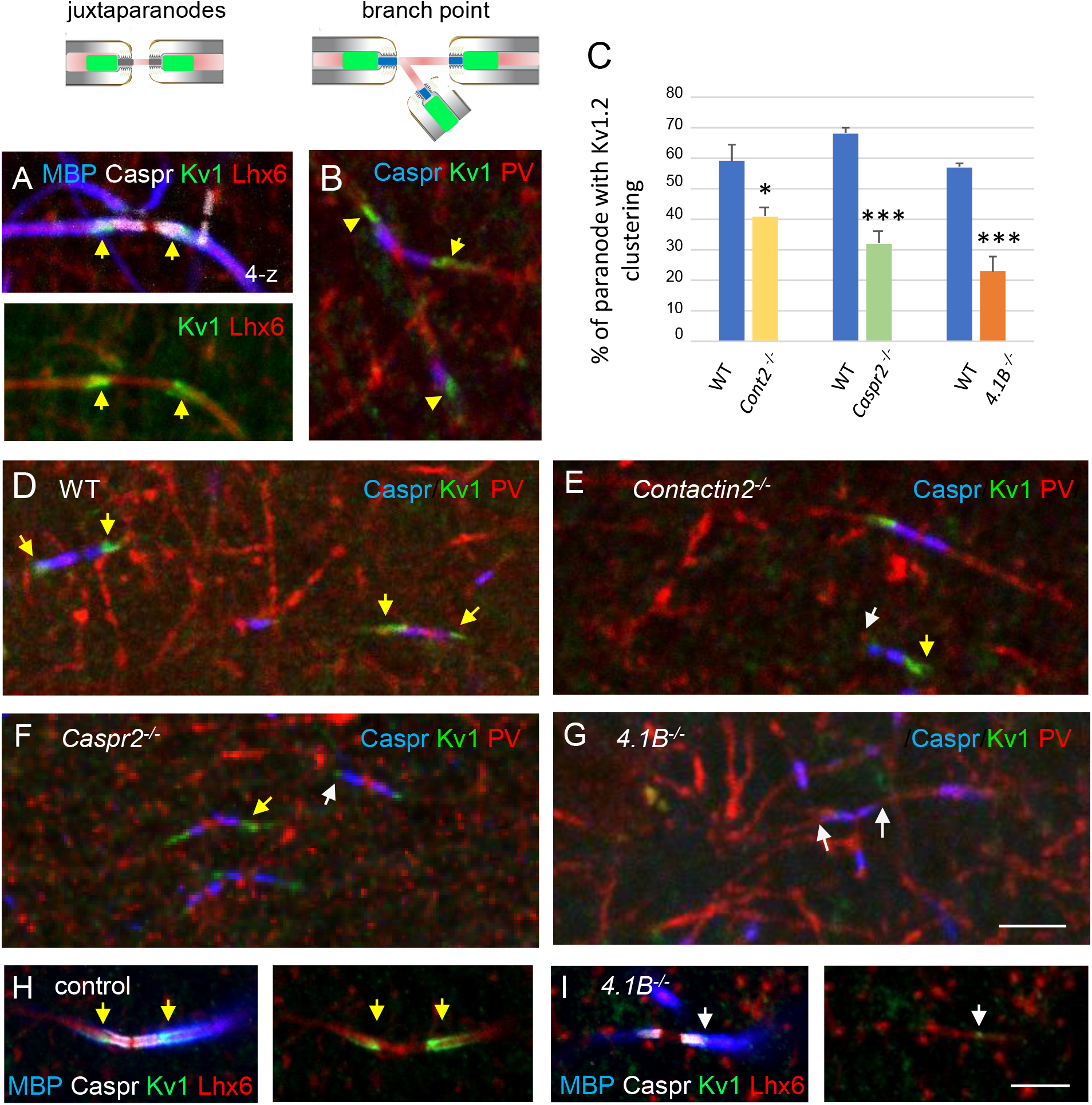
The clustering of juxtaparanodal Kv1 channels is reduced in *Contactin2*^-/-^, *Caspr2*-^-/-^,and *4.1B*^-/-^ GABAergic axons. Immunostaining for Kv1.2 channels in GABAergic axons of the stratum radiatum in CA1 at P70. A, H: Juxtaparanodal clustering of Kv1 channels in *Lhx6-Cre;tdTomato* control mouse immunostained for MBP (blue), Caspr (white) and Kv1.2 (green). tdTomato expressed in Lhx6-positive cells is in red. B: Juxtaparanodal clustering of Kv1 channels (green) in wild-type at PV axon branch point (red) with paranodes stained for Caspr (blue). C: Quantification of the percentage of paranodes associated with Kv1.2 clustering in the PV axons of the CA1 stratum radiatum. Mutant mice are compared with their respective controls. Means ± SEM of 7-11 ROls from 3 mice/genotype. Significant difference by comparison with wild-type: * P<0.05; *** P<0.001 using the Mann-Whitney test. D-G: Immunostaining for Kv1.2 (green) in wild-type (D), *Contactin2*^-/-^ (E), *Caspr2*^-/-^ (F), and *4.1B*^-/-^ (G) in PV axons (red) with paranodes stained for Caspr (blue). Yellow arrows point to juxtaparanodal clustering of Kv1 in control or mutant mice and white arrows indicate the lack of proper Kv1 clustering in mutants. Alteration of Kv1 clustering is also observed in *4.1B*^-/-^; *Lhx6-Cre;tdTomato* mice (I compared to control in H). Bar: 5 μm.

**Figure 7:**
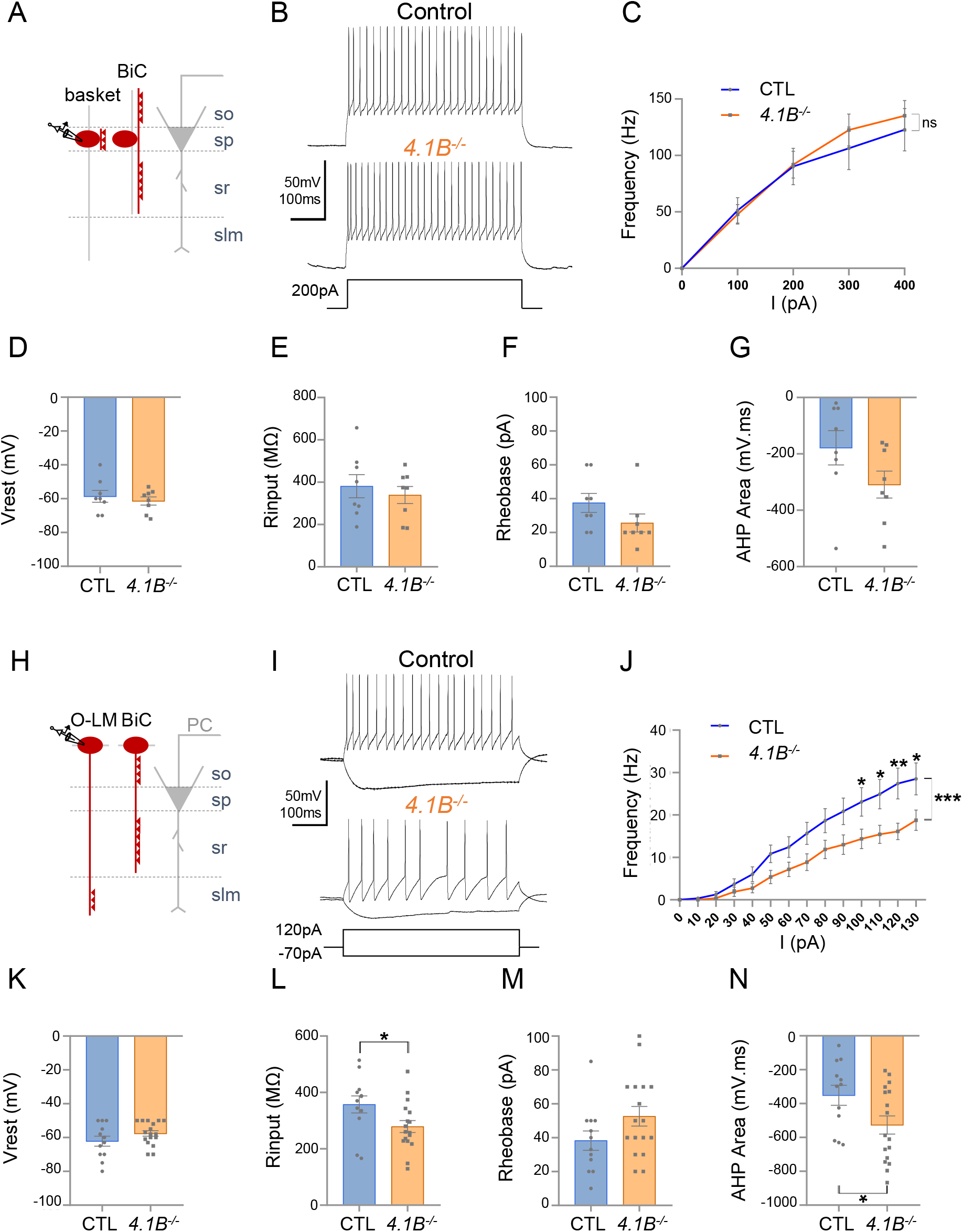
The excitability of O-LM interneurons in the stratum oriens is selectively decreased in *4.1B*^-/-^mice. A-G: Patch-clamp recordings of Lhx6-positive PV fast-spiking interneurons in the stratum pyramidale (SP). A: Schematic illustration of basket and bistratified (BiC) cells in CA1 located in the SP and innervating the soma and dendrites of a pyramidal cell (PC). B: Typical fast-spiking activity evoked by a 200 pA stimulation during 500 ms showing similar firing patterns of 4.1B-deficient and control basket cells. C: *F-I* relationship of mean spike frequency depending on current intensities is not different between the genotypes. The resting membrane potential (Vrest) (D), input resistance (Rinput) (E), rheobase (F) and afterhyperpolarization (AHP) area (G) are not affected in *4.1B*^-/-^ mice. Means ± SEM. Detailed statistical analysis using Mann-Whitney test, see Table 1. H-N: Patch-clamp recordings of Lhx6-positive SST interneurons in the stratum oriens (SO) from control and *4.1B*^-/-^ mice. H: O-LM and bistratified cells in CA1 located in the SO and innervating PC dendrites. I: Example of continuous spike train discharge elicited by a 120 pA depolarizing current pulse during 500 ms and voltage response to a −70 pA hyperpolarizing or in control and *4.1B*^-/-^ mice. J: *F-I* curves. The mean frequency is significantly reduced in *4.1B*-deficient inhibitory neurons compared to controls (two-way ANOVA, ***P<0.0001, **P<0.01, *P<0.05). K: The resting membrane potential is not affected. L: The Rinput is significantly decreased and the rheobase (M) is not changed. N: The AHP area of individual AP is significantly increased (*P<0.05, Mann-Whitney test). For a detailed statistical summary of intrinsic parameters, see Table 2.

**Table 1.** Intrinsic electrophysiological properties of fast-spiking Lhx6-interneurons in the stratum pyramidale

|  | Control (mean±SEM, n=8) | 4.1B<sup>-/-</sup> (mean±SEM, n=8) | MW test, *p* value |
| --- | --- | --- | --- |
| **Passive properties** |  |  |  |
| Resting membrane potential (mV) | -58.63±3.53 | -61.38±2.40 | MW, 0.7391 |
| Input resistance (MΩ) | 381.13±54.47 | 339.25±40.15 | MW, 0.5737 |
| Rheobase (pA) | 37.40±5.59 | 25.63±5.30 | MW, 0.0998 |
| **Action potential parameters** |  |  |  |
| Amplitude (mV) | 59.33±3.73 | 55.22±2.51 | MW, 0.4256 |
| Area (mV.s) | 60.64±6.43 | 58.34±5.19 | MW, 0.3823 |
| Half-width (ms) | 0.95±0.08 | 0.98±0.07 | MW, 0.9397 |
| Threshold (mV) | -46.46±2.56 | -46.45±1.97 | MW, >0.9999 |
| AHP area (mV.ms) | -178.40±60.73 | -309.07±47.69 | MW, 0.1304 |
| AHP peak (mV) | -9.06±1.50 | -9.73±0.83 | MW, 0.5737 |

Statistical differences between groups were calculated with a Mann-Whitney test (MW). Control: n=8 mice; 4.1B<sup>-/-</sup>: n=5 mice.

**Table 2.** Intrinsic electrophysiological properties of Lhx6-interneurons in the stratum oriens

|  | Control (mean±SEM, n=12) | 4.1B<sup>-/-</sup> (mean±SEM, n=17) | MW test, *p* value |
| --- | --- | --- | --- |
| **Passive properties** |  |  |  |
| Resting membrane potential (mV) | -62.17±2.93 | -57.59±1.65 | MW, 0.2527 |
| Input resistance (MΩ) | 357.33±30.30 | 278.94±21.12 | MW, 0.0236* |
| Rheobase (pA) | 38.25±5.73 | 52.65±5.79 | MW, 0.1221 |
| **Action potential parameters** |  |  |  |
| Amplitude (mV) | 65.34±1.84 | 64.06±1.94 | MW, 0.6788 |
| Area (mV.s) | 111.44±17.87 | 104.80±8.93 | MW, 0.8788 |
| Half-width (ms) | 1.61±0.25 | 1.63±0.16 | MW, 0.5929 |
| Threshold (mV) | -45.69±1.83 | -43.65±1.43 | MW, 0.5558 |
| AHP area (mV.ms) | -351.29±58.99 | -526.59±53.50 | MW, 0.0380* |
| AHP peak (mV) | -12.57±1.79 | -16.24±1.25 | MW, 0.1280 |

Statistical differences between groups were calculated with a Mann-Whitney test (MW). Control: n=9 mice; 4.1B<sup>-/-</sup>: n=8 mice.

Next, we asked whether such myelin loss could be due to a reduced number of oligodendrocytes in the hippocampus using immunostaining for Olig2 as a marker for the oligodendrocyte lineage (Fig. 3A-D). The density of Olig2-positive cells in the stratum radiatum was 22.3 ± 1.6/10^5^ μm^2^ in the 4.1B KO mice versus 25.3 ± 2.2/10^5^ μm^2^ in wild-type animals at P35 (Fig. 3E). We did not observe any decrease in the number of Olig2-positive cells in the CA1 layers that could explain the dramatic loss of myelin in 4.1B KO mice. We also examined the density of Olig2-positive cells in the dentate gyrus since the neuronal progenitors of the granule cell layer might contribute to the generation of oligodendrocytes in demyelinating pathological conditions such as in the cuprizone toxic model (Jessberger et al., 2008; Klein et al., 2020). The pattern of MBP-positive axons was apparently slightly disorganized in the molecular layer and not in the hilus of the dentate gyrus in 4.1B^-/-^ mice at P35 (Fig. 3D). We did not observe any difference in the density of Olig2-positive cells in the hilus, granule cell or molecular layers of the dentate gyrus (Fig. 3F). These results suggest that the developmental alteration of myelin induced by the loss of the axonal cytoskeletal linker 4.1B may not be associated with change in oligodendrogenesis.

### 4.1B-deficient GABAergic axons display reduced internodal length

Protein 4.1B mediates interactions between membrane adhesion complexes and the spectrin cytoskeleton and binds Caspr at paranodes as schematized in Fig. 4A. We previously showed that 4.1B promotes via its interaction with Caspr, the growth of oligodendroglial processes and myelin sheath convergence in the developing spinal cord (Brivio et al., 2017). Immunofluorescence staining for 4.1B in the hippocampus indicated that the scaffolding protein was expressed by GABAergic PV interneurons and not by pyramidal neurons in the CA1 pyramidal layer at P70 (Fig. 4C). As a control for specificity, no staining was observed on hippocampal sections from *4.1B*^-/-^ mice (Fig. 4D). Protein 4.1B immunostaining was present around the soma of PV interneurons, along the internodes and enriched at paranodes whereas the nodal gap was unlabeled (Fig. 4E, E’). We asked whether the elongation of internodes could be affected in the 4.1B-deficient CA1 hippocampus at P70 and analyzed the distribution of the length of internodes (Fig. 4B). Analysis of the cumulative frequency indicates a significant difference between the genotypes (P<0.0001; Kolmogorov-Smirnov test). The mean internodal length was significantly reduced (P=0.0286, Mann-Whitney test) in 4.1B KO (34.8 ± 2.9 μm; n=191) by comparison with wild-type (43.8 ± 0.3 μm; n= 275) mice. The maximum length was similarly decreased to 93.8 μm in the 4.1B KO versus 114.5 μm in the wild-type mice. The reduction of internodal length strongly indicates that protein 4.1B may be implicated in myelin sheath elongation.

### 4.1B-deficiency causes misdistribution of nodal and heminodal Nav channels in hippocampal GABAergic interneurons

At the paranodal junction, the axonal cytoskeleton including 4.1B and ßII-spectrin also organizes the boundary between the internodes and the nodal Nav channels (Fig. 4A) (Brivio et al., 2017; Buttermore et al., 2011; Zhang et al., 2013; Zonta et al., 2008). Accordingly, the Nav complex is mislocalized distant from the paranodes in the developing spinal cord of the 4.1B KO mice before the convergence of internodes (Brivio et al., 2017). In the adult spinal cord, this phenotype is no longer observed and the nodal Nav clusters become properly juxtaposed to the paranodes in mature nodes. Hippocampal inhibitory axons are covered by discontinuous myelin with persistence of heminodal structures in the adult, and myelination of ramified axons implies the presence of nodes at branch points so that it was particularly interesting to analyze whether the distribution of Nav channels may be disturbed in the 4.1B-deficient axons.

We first analyzed the architecture of the nodes of Ranvier in the GABAergic interneurons of the CA1 hippocampus of wild-type mice at P30 and P70 by performing panNav channel immunostaining. As illustrated in Fig. 5A, myelination along Lhx6-positive axons was partly discontinuous in the stratum radiatum at P70, leaving uncovered axonal segments (asterisks). Nav channels were detected at nodes (Fig. 5B, B’, G) and branch points (Fig. 5C, C’). Heminodal clustering of Nav channels was observed for interspaced internodes (Fig. 5E). We noticed that the lengths of nodal Nav immunostaining measured between two paranodes were higher at P30 than at P70. Analysis of the cumulative frequency indicates that the distribution of nodal Nav immunostaining lengths was significantly different with age (P=0.0264; Kolmogorov-Smirnov test). At the age of P30, 44% of the nodes displayed a length > 2 μm versus only 22 % at P70, indicating that the internodes may still elongate during the late developmental stage (Fig. 5D).

We then investigated whether 4.1B deficiency may induce alteration of the distribution of nodal and heminodal Nav channels in the CA1 hippocampus at P70. In 4.1B-deficient mice, a number of inhibitory PV or SST axons display interrupted myelination when entering the stratum radiatum showing heminodes (Fig. 2H, J and S4B). Interestingly, we observed at heminodes of 4.1B-deficient inhibitory axons, a gap between the site of Nav clustering and the paranode stained for Caspr (Fig. 5F and Fig. S4B) as compared to control mice (Fig. 5E and S4A). The gap could reach 10 μm and displayed a mean value of 2.25 ± 0.51 μm significantly different from the control value (0.04 ± 0.02 μm) (n=25 heminodes, 2 mice/genotype; P<0.0001, Mann-Whitney test) (Fig. 5I). By contrast, the length of heminodal Nav clusters was similar in control (1.74 ± 0.19 μm) and in 4.1B KO (1.84 ± 0.12 μm) mice (P=0.427, Mann-Whitney test) (Fig. 5J). A gap could also be observed between nodal Nav and convergent internodes stained for MBP (Fig. 5H) in 4.1B-deficient mice by comparison with control ones (P=0.0029, Mann-Whitney test) (Fig. 5I), together with an increase in the length of nodal Nav clusters (Fig. 5J). The mean length of nodal Nav in 4.1B KO (1.92 ± 0.25 μm, n=26) was significantly increased by comparison with control mice (1.59 ± 0.14 μm, n=87) (P=0.0068, Mann-Whitney test). Thus, the positioning of Nav clusters is significantly disturbed at the nodes and heminodes of dysmyelinated GABAergic interneurons of 4.1B-deficient hippocampus.

### Juxtaparanodal alterations in hippocampal interneurons of Contactin2-, Caspr2- and 4.1B null mutant mice

Finally, we analyzed the distribution of the juxtaparanodal Kv1 channels in the stratum radiatum at P70. As expected, immunostaining for Kv1.2 was detected at the juxtaparanodes or hemi-juxtaparanodes under the compact myelin including at branch points both in Lhx6- and PV-positive axons (Fig. 6A, B). We investigated whether the deficiency in Kv1-associated CAMs may induce alteration of the juxtaparanodes in hippocampal interneurons like it was observed in myelinated tracts of the corpus callosum, optic nerves, or spinal cord for Contactin2 KO and Caspr2 KO mice (Poliak et al., 2003; Savvaki et al., 2008; Traka et al., 2003). As analyzed in double-blind experiments, Contactin2- or Caspr2-deficiency induced a strong decrease in Kv1 channel clustering at the juxtaparanodes of PV myelinated axons in the CA1 hippocampus at P70 (Fig. 6D-F) although a residual expression of Kv1 was still present at some juxtaparanodes in both mutants. The percentage of paranodes bordered by Kv1.2 immunostaining in PV axons was significantly reduced by 30% in Contactin2 KO and 53% in Caspr2 KO mice (P= 0.014 and P<0.0001, Mann-Whitney test, respectively) (Fig. 6C). The scaffolding protein 4.1B is known to bind Caspr2 and to be required for the proper assembly of the juxtaparanodes both in the PNS and CNS (Buttermore et al., 2011; Cifuentes-Diaz et al., 2011; Einheber et al., 2013). We observed a dramatic decrease (−61%; P<0,0001, Mann-Whitney test, n=9 ROIs, 3 mice/genotype) in the Kv1 clustering at juxtaparanodes of myelinated PV axons of 4.1B KO mice at P70 (Fig. 6G). *4.1B*^-/-^ mice crossed with the *Lhx6-Cre;tdTomato* line showed similar alteration of Kv1 clustering at juxtaparanodes in Lhx6-positive axons as compared with control mice (Fig. 6H and I). In summary, deficiency in each of the Kv1-associated proteins, Contactin2, Caspr2, and 4.1B induced alteration of juxtaparanodal Kv1 clustering along myelinated GABAergic axons in the hippocampus.

### The excitability of Lhx6-positive interneurons is differently affected in the stratum oriens and stratum pyramidale of 4.1B KO mice

We showed that the 4.1B KO mice exhibited a dramatic and selective loss of myelin coverage of the inhibitory axons crossing the stratum radiatum. In particular the SST O-LM interneurons showed a lack of myelin. The PV axons crossing the stratum radiatum that may belong to bistratified interneurons were also dysmyelinated whereas PV axons of basket cells seemed to be more preserved in the stratum pyramidale. These observations led us to further question whether the phenotypes observed could be associated with physiological changes in hippocampal interneurons. We first recorded the activity of Lhx6Cre;tdTomato-positive cells in the stratum pyramidale of control and 4.1B KO mice (Figure 7A-G). Different subtypes of Lhx6-interneurons are located in the stratum pyramidale as schematized in Fig. 7A, including the fast-spiking basket cells innervating the soma of pyramidal cells and bistratified cells contacting their basal and apical dendrites. We performed whole-cell patch-clamp in current clamp configuration recordings on P30 acute slices and selected fast-spiking interneurons displaying a typical continuous non-stuttering discharge (Hu et al., 2014) with a frequency above 50 Hz under injection of a 200 pA current intensity (Fig. 7B). As shown by the *F-I* curve during injection of current steps of 100 pA increments, the firing frequencies were unchanged in *4.1B*^-/-^ compared to control mice (P=0.5505, two-way ANOVA) (Fig. 7B and C). The resting membrane potential (Vrest) (−61.38 ± 2.40 mV in *4.1B*^-/-^ vs - 58.63 ± 3.53 mV in controls, P=0.7391 using Mann-Whitney test) and the input resistance (Rinput) (339.25 ± 40.15 MΩ in *4.1B*^-/-^ vs 381.13 ± 54.47 MΩ in controls, P=0.5737 using Mann-Whitney test) were not changed (Fig. 7D and E). Similarly, the rheobase (25.63 ± 5.30 pA in *4.1B*^-/-^ vs 37.40 ± 5.59 pA in controls, P=0.0998, Mann-Whitney test) (Fig. 7F) and afterhyperpolarization (AHP) area (−309.07 ± 47.69 mV.ms in *4.1B*^-/-^ vs −178.40 ± 60.73 mV.ms in controls, P=0.1304, Mann-Whitney test) were not significantly affected (Fig. 7G). Overall, we found no significant modification in the intrinsic electrophysiological properties of fast-spiking basket cells (Table 1). Thus, the PV basket cells, which axons only displayed mild alteration of myelin in the stratum pyramidale, did no exhibit alteration of their excitability.

To investigate the electrophysiological properties of O-LM cells, we next recorded Lhx6Cre;tdTomato-positive interneurons in the stratum oriens. As shown in the schematic illustration, different subtypes of Lhx6-interneurons are located in the stratum oriens including SST O-LM cells innervating distal dendrites and bistratified cells connecting basal and apical dendrites of pyramidal cells (Fig. 7H). The O-LM cells are the most abundant subtype of Lhx6-interneurons in the stratum oriens and display continuous discharge activity in contrast to the bistratified cells (Tricoire et al., 2011). Here, we selected Lhx6-positive O-LM cells that displayed a non-stuttering and continuous pattern of discharge during spike trains (500 ms) as illustrated by traces for control and *4.1B*^-/-^ mice upon 120 pA current injection (Fig. 7I). We observed a marked decrease in the mean discharge frequency of *4.1B*^-/-^ cells as indicated by the *F-I* curve depending on injected current steps of 10 pA increments (P<0.0001, two-way ANOVA). Notably, the discharge frequency was significantly decreased upon 100, 110, 120 and 130 pA stimulation (P=0.037, P=0.018, P=0.002, P=0.014, respectively; two-way ANOVA and Sidack post-test) (Fig. 7J). The resting membrane potential (Vrest) was unchanged (−57.59 ± 1.65 mV in *4.1B*^-/-^ vs −62. 17 ± 2.93 mV in controls, P=0.2527, Mann-Whitney test) (Fig. 7K). Hyperpolarizing current steps were injected for analyzing input resistance (Rinput) that was significantly decreased by 22% (278.94 ± 21.12 MΩ in *4.1B*^-/-^ vs 357.33 ± 30.30 MΩ in controls, P=0.0236, Mann-Whitney test) (Fig. 7L). The rheobase was not significantly changed (52.65 ± 5.79 pA in *4.1B*^-/-^ vs 38.25 ± 5.73 pA in controls, P=0.1221, Mann-Whitney test) (Fig. 7M). The intrinsic parameters of APs were also analyzed showing an increased AHP area (−526.59 ± 53.50 mV.ms in *4.1B*^-/-^ vs −351.29 ± 58.99 mV.ms in controls, P=0.0380, Mann-Whitney test) (Fig. 7I, N and Table 2). These data indicated that the dysmyelinating phenotype of *4.1B*^-/-^ mice in O-LM cells was associated with a selective decrease in their excitability.

### Structural modification of the Axon Initial Segment of O-LM interneurons in *4.1B*^-/-^ mice

The efficacy of spike generation is related to the structural organization of AIS, which could be modified in dysmyelinated axons. We investigated whether the decreased excitability in SST O-LM cells may be due to differences in the localization of AP initiation site. We first used the method of multiple derivatives of the somatic AP signal that allows to visualize a separation of the axonal and somatodendritic APs (Fig. 8A and B). Our results showed that 58% of control stratum oriens interneurons including O-LM cells had a stationary inflection in the rising phase of the AP (Fig. 8A2 and C top pie chart). We calculated the second (d^2^V/dt^2^) and third (d^3^V/dt^3^) derivatives to better illustrate separation of two inflection points (Fig. 8A3, arrow in A4) as an indication of the distant site of the AP initiation from the soma. In contrast, 64% of *4.1B*^-/-^ interneurons did not display stationary inflection (Fig. 8B2-B4 and C bottom pie chart). This statistically significant difference (P=0.0017, Fisher’s exact test) suggests that the AP was initiated closer to the soma in *4.1B*^-/-^ neurons than in control ones. This could be due to a modification of AIS geometry such as a shortening or a displacement relative to the soma.

**Figure 8:**
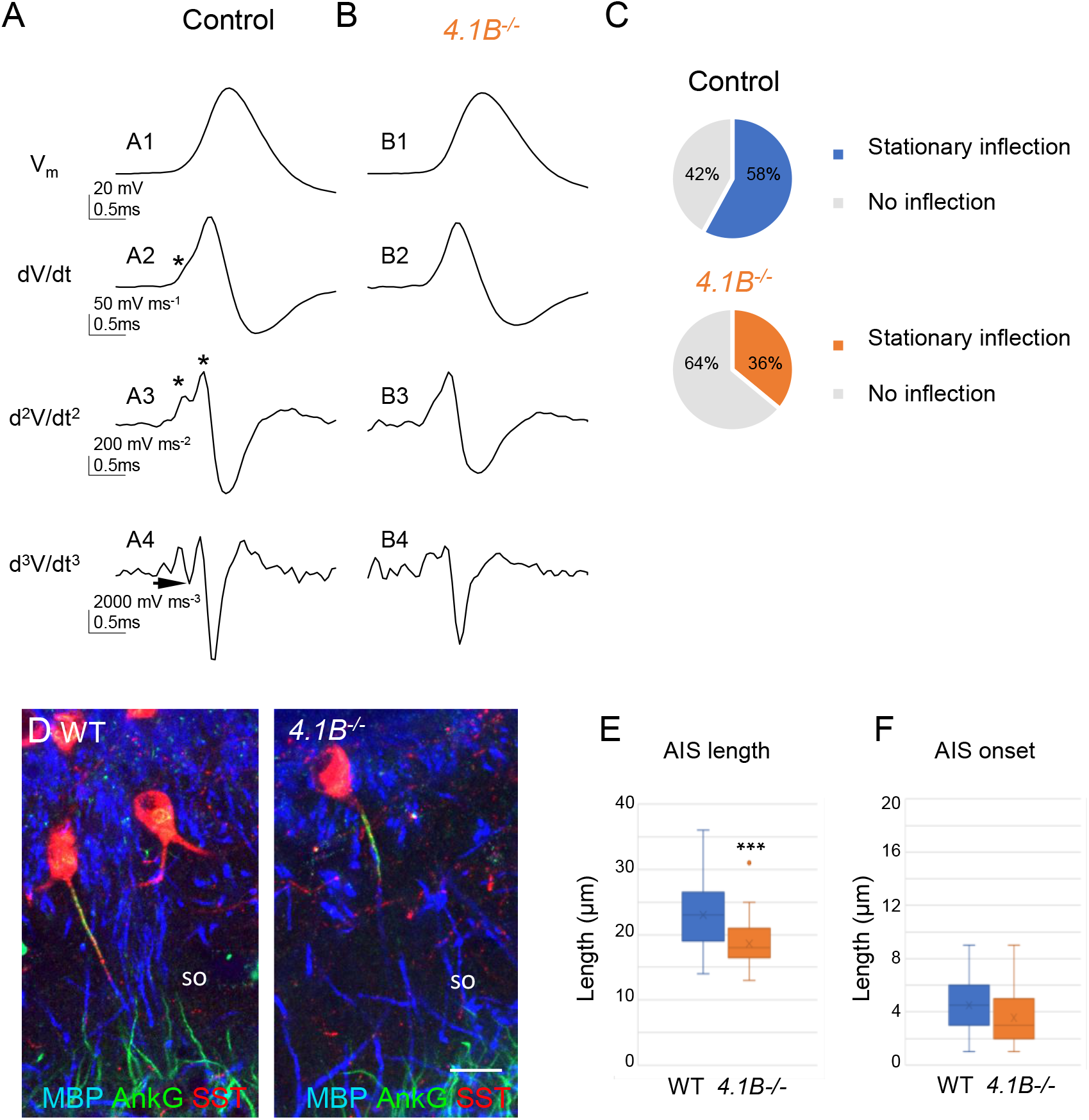
Structural modification of the AIS of O-LM interneurons in 4.1B^-/-^mice. A-C: Analysis of somatic action potentials (APs). Representative examples illustrating the voltage waveform of the AP (A1 and B1; in mV) from a control (A1) and a *4.1B*^-/-^ neurons (B1). Below are presented the first (A2 and B2; dV/dt, in mV/ms), second (A3 and B3; d^2^V/dt^2^, in mV/ms^2^), and third derivatives (A4 and B4; d^3^V/dt^3^, in mV/ms^3^). The stationary inflection in the d^2^V/dt^2^ trace is indicated with asterisks. The third derivative allows to detect the second inflection point when the trace reaches or goes under zero (arrow). C: We observed a second inflection point in 58% of control (top pie graph) and 36% of *4.1B*^-/-^ cells (bottom pie graph). This difference in the number of cells displaying an inflection point (asterisks in A2 and A3) is significant (P=0.017, Fisher’s exact test). This could be due to the AIS shortening in *4.1B*^-/-^ O-LM interneurons. D: Hippocampal sections of wild-type and *4.1B*^-/-^ mice at P35 immunostained for MBP (blue), SST (red) and AnkyrinG (green). E: Length of AIS measured for AnkyrinG immunostaining of SST interneurons in the stratum oriens (SO). Significant difference by comparison with wild-type: *** P<0.001 using Mann-Whitney test. F: Distance of AIS onset from the soma is not changed between wild-type and *4.1B*^-/-^ mice (P=0.099, Mann-Whitney test; n=22 in wild-type and n=32 cells in *4.1B*^-/-^ mice; 3 mice/genotype). Bar: 10 μm in D.

We thus evaluated whether the loss of myelin in hippocampal O-LM interneurons could be associated with remodeling of the AIS using immunostaining for AnkyrinG. We observed that the AIS of SST interneurons in the stratum oriens was significantly shorter (Fig. 8D, E) in 4.1B KO by comparison with wild-type mice (18.6 ± 0.7 μm, n=29 in *4.1B*^-/-^ vs 23.0 ± 0.9 μm, n=29 in wild-type; P=0.0004, Mann-Whitney test). The AIS onset measured from the soma or the primary dendrite was not modified (3.8 ± 0.5 μm in *4.1B*^-/-^ vs 4.8 ± 0.5 μm in wild-type; P=0.099, Mann-Whitney test) (Fig. 8F). Taken together, our results indicated that dysmyelination in *4.1B*^-/-^ mice caused a shortening of AIS and a net decrease in the intrinsic excitability of O-LM cells.

### Inhibitory inputs onto CA1 pyramidal neurons are influenced by dysmyelinated *4.1B*^-/-^ phenotype

We then asked the question of the role of interneuron myelination in the proper inhibitory synaptic transmission on pyramidal cells. First, we recorded spontaneous inhibitory postsynaptic currents (sIPSCs) on pyramidal cells in APV (40 μM) and CNQX (10 μM) condition (Figure 9A and B). We showed a significant increase in sIPSCs peak amplitude in the *4.1B* KO mice (45.61 ± 2.93 pA in *4.1B*^-/-^ vs 31.51 ± 2.93 pA in controls; P=0.0022, Mann-Whitney test). The frequency of sIPSCs also tended to be increased (4.97 ± 1.02 Hz in *4.1B*^-/-^ vs 2.76 ± 1.13 Hz in controls, P=0.1320, Mann-Whitney test) (Fig. 9C). Interestingly, the occurrence probability of small amplitude sIPSCs was decreased in *4.1B* KO mice (P<0.0001, Kolmogorov-Smirnov test) (Fig. 9D). To further examine the properties of inhibitory synaptic inputs, we recorded miniature inhibitory postsynaptic currents (mIPSCs) on pyramidal cells as AP-independent inhibition. We found that neither the mean amplitude (28.29 ± 2.33 pA in *4.1B*^-/-^ mice vs 24.57 ± 2.68 pA in controls, P=0.2224, Mann-Whitney test) nor the mean frequency were affected in *4.1B*^-/-^ mice (2.41 ± 0.47 Hz in *4.1B*^-/-^ mice vs 2.73 ± 0.51 Hz in controls, P=0.5457, Mann-Whitney test). This data on mIPSCs indicated that the *4.1B*^-/-^ mutation may not modify the inhibitory pre-synaptic terminals connecting pyramidal cells.

**Figure 9:**
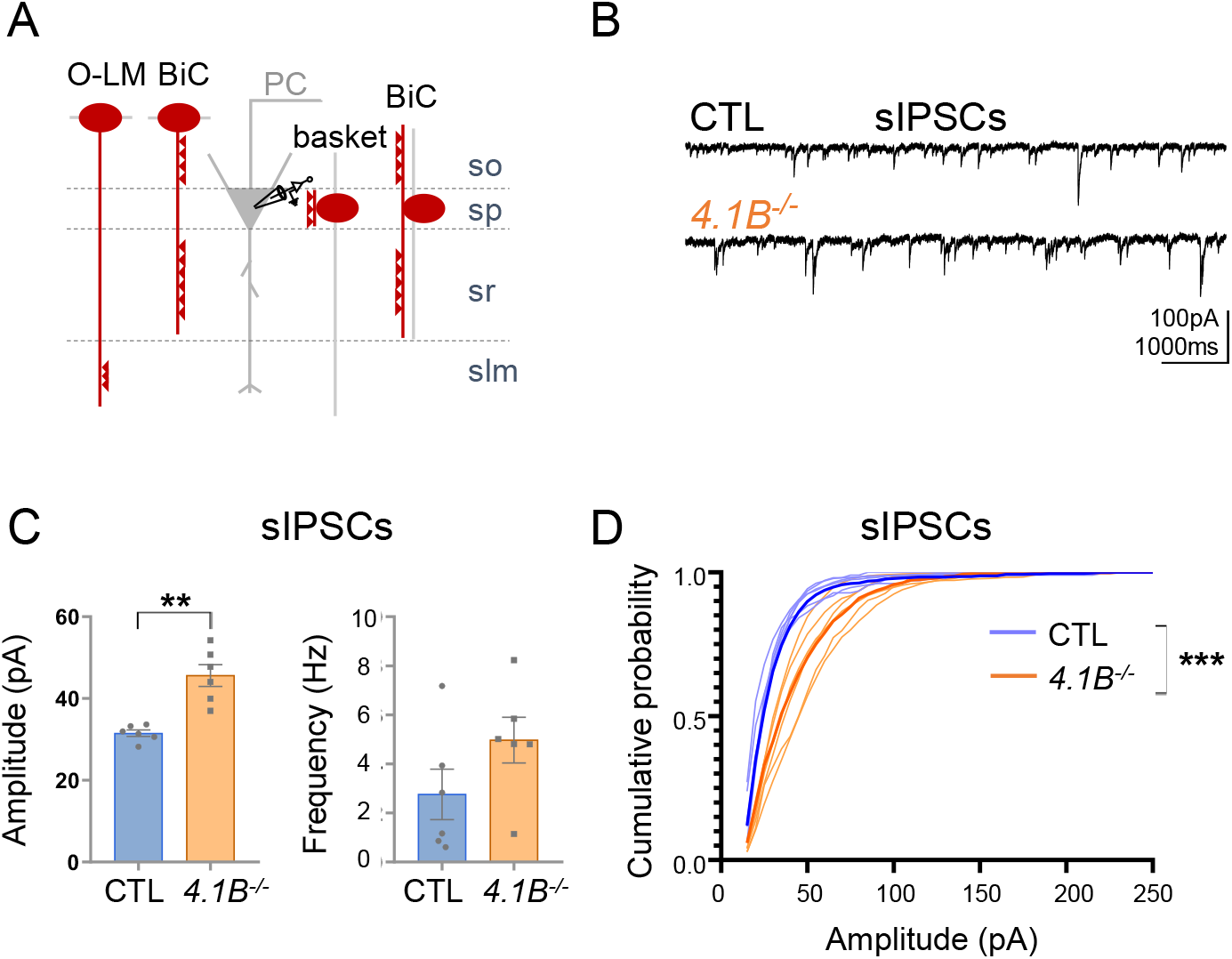
Inhibitory inputs onto CA1 pyramidal neurons are affected in *4.1B*^-/-^ mice. A: Schematic illustration of inhibitory inputs onto the soma, proximal and distal dendrites of pyramidal cells. B: Example traces of sIPSCs onto CA1 pyramidal neurons of control and *4.1B*^-/-^ mice. C: The amplitude of sIPSCs is significantly increased in *4.1B*^-/-^ mice (**P=0.0022) and the frequency is not affected (P=0.1320); 15 to 400 pA events analyzed during 3 min. Statistical values were performed with a Mann-Whitney test. D: Cumulative probability for the amplitude of sIPSCs (3 min, 6 cells and 4-5 mice/genotype) showing a significant different distribution between genotypes (Kolmogorov-Smirnov test, ***P<0.0001). Note that the probability of small amplitude events is reduced in *4.1B*^-/-^ mice suggesting that distal dendritic inhibition by O-LM cells could be altered.

The increased amplitude of sIPSCs in *4.1B*^-/-^ mice could reflect an enhanced proximal and a reduced distal GABAergic inhibition onto pyramidal cells. Taken together, these results suggest that the distal inhibitory inputs onto pyramidal cells may be reduced as a consequence of the dysmyelination and decreased excitability of O-LM cells.

### Spatial memory is altered in *4.1B*^-/-^ mice

The CA1 inhibitory interneurons have been implicated in spatial learning and memory (Jeong and Singer, 2022). We tested whether the severe loss of myelin in interneurons may induce alteration of hippocampus-based place learning and memory using the Y-maze and Barnes maze tests. The spatial working memory was affected in *4.1B*^-/-^ mice as tested through the Y-maze (Fig. 10A). Spontaneous alternation was measured during 8 min and the percentage of alternation was significantly decreased in 4.1B KO (53.5 ± 2.9 %) versus control (64.0 ± 2.1) mice (n=8 per genotype, P=0.0068, Student’s t test). The number of total entries in maze arms did not differ between the genotypes (33.6 ± 3.3 in 4.1B KO vs 35.1 ± 4.0 in control mice, P=0.7623, Student’s t test) as an indication that the locomotor activity is not affected. Next, we analyzed the Barnes maze task performance. Acquisition learning trials were performed during 4 days (2 trials/day) and the probe test the day after during which the escape box was removed. Decrease in latency to locate target was observed during training in each genotype as an indication that spatial learning occurred in *4.1B*^-/-^ mice. We observed that the latency to target or the latency to enter the escape box was significantly increased in the 4.1B KO mice by comparison with controls (two-way ANOVA, genotype effect: P=0.0157 and P=0.0153, respectively; WT n=14; *4.1B*^-/-^n=11) (Fig. 10B). Since the simple quantification of latency describing performance in the Barnes maze may result in only a partial understanding of mouse behavior and cognitive capacity, we analyzed and classified exploration paths into five strategies including spatial (direct, corrected, long-correction) and non-spatial (serial or random) search strategies as illustrated in Fig. 10C, D, F, G) (Cheng et al., 2019; Illouz et al., 2016). The *4.1B*^-/-^ mice appeared to adopt distinct search strategies by comparison with control mice during the learning trials. Control mice used spatial (57%), serial (29%) or random (14%) strategies during the first acquisition day of training. The frequency of the spatial search strategy increased to 86% for the probe test with only 14% of mice using of a serial approach (Fig. 10E). In contrast, in 4.1B KO mice, the random approach was by far the most dominant strategy (73 %) on the first acquisition day, the spatial search strategy being used 18% of the time. There was an effective evolution of the *4.1B*^-/-^ mice to use most frequently a spatial search strategy (64%) for the probe test, whereas 9% used serial and 27% random exploration paths (Fig. 10H). Scaling of search strategies from direct=1 (best strategy) to random=0 (worst strategy) gives a cognitive score that was significantly (P<0.0001, two-way ANOVA) reduced in the 4.1B KO mice (Fig. 10I). In conclusion, the selective dysmyelination of hippocampal interneurons was associated with impairment of working and spatial memories and cognitive capacity in the 4.1B-deficient mice.

**Figure 10:**
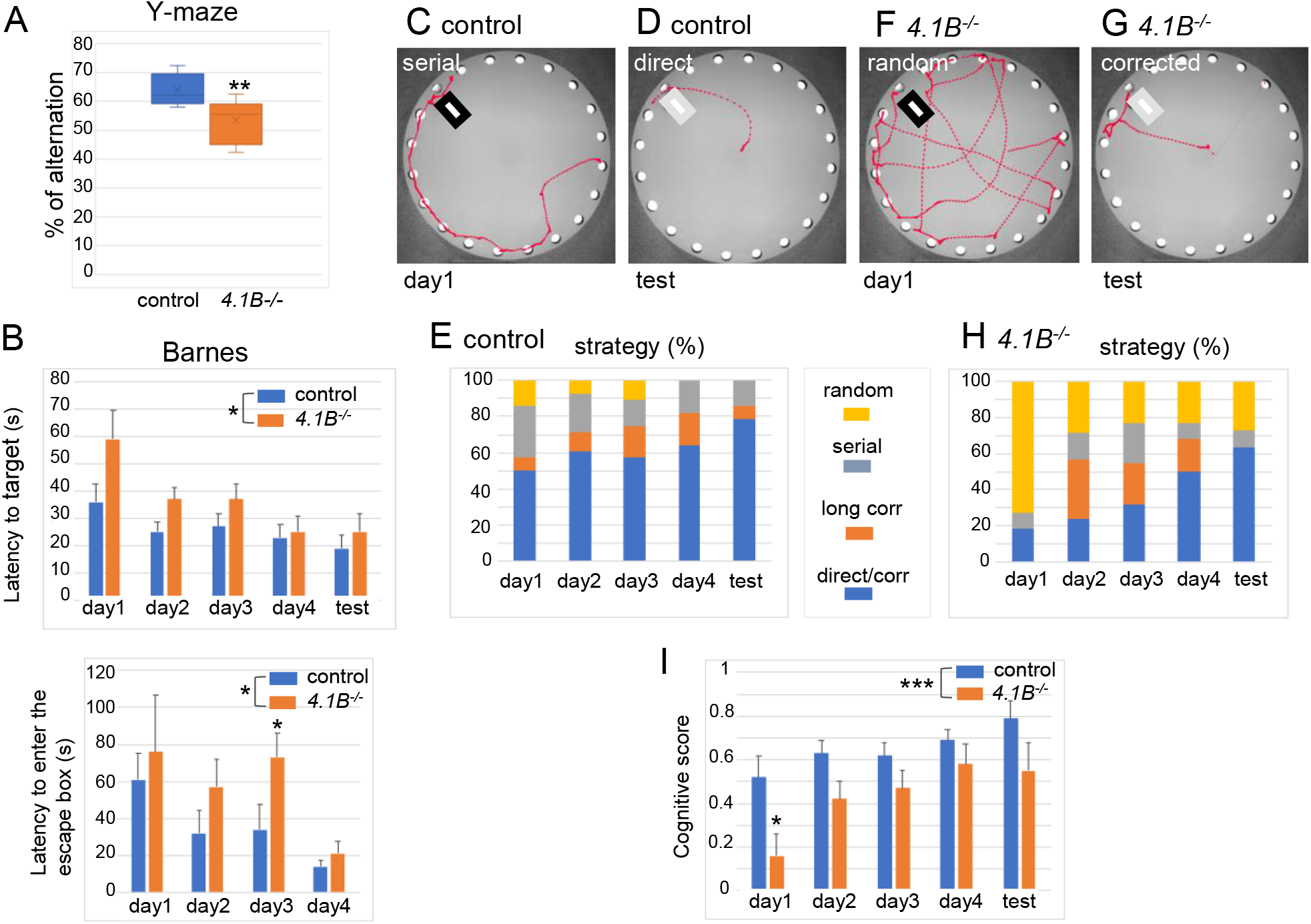
Alteration of working and spatial memory in 4.1B-deficient mice. Control and *4.1B*^-/-^ mice with *Lhx6-Cre;tdTomato* genotype were tested for Y-maze and Barnes maze. A: Spontaneous alternation in Y-maze was measured during 8 min and the percentage of alternation was significantly decreased in 4.1B-deficient mice (n=8 per genotype, P<0.01 using the Student’s t-test). B: Barnes maze task performance. Acquisition learning trials were performed during 4 days (2 trials/day) and the probe test the day after during which the escape box was removed. The latency to locate or to enter the escape box was noted and the mean performance of two trials was expressed as mean ± SEM. C-I: Exploration paths were analyzed and classified into five strategies including spatial (direct, corrected, long-correction) and non-spatial (serial or random) search strategies. Examples of serial (C) or random (F) on day 1 and direct (D) or corrected (G) strategies during the probe test are provided. The percentage of each strategies is shown for control (E) and *4.1B*^-/-^ mice (H). I: Strategies were scaled from direct=1 (best strategy), corrected=0.75, long-correction=0.5, serial=0.25 and random=0 (worst strategy) to give a cognitive score for each trial. Significant difference of *4.1B*^-/-^ mice by comparison with controls using two-ways ANOVA, * P<0.05 and **** P<0.0001; controls n=14, *4.1B*^-/-^ n=11).

## Discussion

Our results describe for the first time a genetic model of selective myelin loss in the hippocampus. We show that the extent of myelination of PV and SST axons is severely reduced in the stratum radiatum. Deficiency in the scaffolding protein 4.1B also impairs Kv1 clustering at juxtaparanodes and proper localization of Nav at the nodes of Ranvier. The excitability of the O-LM cells in the stratum oriens is decreased while the properties of the fast-spiking interneurons in the stratum pyramidale are unchanged. Our data suggest that this excitability defect reduces the distal inhibitory drive onto pyramidal cells. Last but not least, behavioral tests indicate that alteration of spatial memory and cognitive performance are related phenotypes to the *4.1B*^-/-^ genotype.

### 4.1B is implicated in the myelin coverage of hippocampal inhibitory axons

Despite the well-established physiological importance of myelination to speed axonal conduction, relatively little is known about its significance for GABAergic local interneurons. Recent reports have highlighted the fact that the different subtypes of GABAergic neurons could be myelinated and suggested that oligodendrocytes can recognize the class identity of individual types of interneurons that they target (Zonouzi et al., 2019). PV- and to a lesser extent SST-expressing cells in the CA1 mouse hippocampus are frequently myelinated on their proximal axonal segments, independently of their subtype identity (Micheva et al., 2016; Stedehouder et al., 2017) The sparse myelination of inhibitory ramified axons implies the presence of heminodes along mature axons. One inherent feature that limits myelin sheath growth and determines node position is the branching pattern. An intriguing question is whether axonal cues may be instructive to determine the extent of myelin sheath coverage. It is known that hippocampal inhibitory axons can form pre-nodal clusters of Nav channels along premyelinated axons (Bonetto et al., 2019; Freeman et al., 2015). At a later stage of development, such prenodes of Nav/Neurofascin/AnkyrinG may indicate positions for limiting myelin sheath extension (Malavasi et al., 2021; Thetiot et al., 2020; Vagionitis et al., 2022). During development, components of the nodes of Ranvier accumulate adjacent to the ends of myelin sheaths to form heminodes prior to forming nodes. The developing myelin sheaths may stop growing when they meet a prenodal cluster, which would be instructive for the position of the node. Indeed, mutation of the nodal CAM Neurofascin-186 has been reported to increase internodal distance (Vagionitis et al., 2022). Here, we observed that deletion of 4.1B reduces the mean internodal length indicating that myelin sheath elongation is restricted in the mutant mice. 4.1B-deficiency strongly reduces myelin ensheathment of both PV and SST axons in the stratum radiatum, by 58 and 63%, respectively. The scaffolding 4.1B, which mediates the linkage between Caspr and the ßII-spectrin may be implicated as a paranodal cue modulating the myelin coverage of GABAergic axons. The association of 4.1 proteins with their transmembrane receptors and the actin-spectrin cytoskeleton can be negatively modulated by phosphorylation (Wang et al., 2014) so that 4.1B may act as a clutch to promote myelin sheath elongation depending on axonal signaling. Whether the paranodal cytoskeleton including 4.1B may be implicated in adaptive myelination deserves further investigation.

The localized effect of 4.1B-deficiency on the myelination of hippocampal interneurons is intriguing. The stratified organization of the hippocampus likely reveals the selective role of 4.1B in promoting myelin sheath elongation in inhibitory interneurons. In the stratum radiatum, we evaluated that as much as 75 % of the myelinated axons are Lhx6-positive. This is in accordance with other studies indicating that the fraction of myelin ensheathing GABAergic neurons is reaching up 80 % in the stratum pyramidale and stratum radiatum (Stedehouder et al. 2017). We did not detect any dramatic loss of myelin immunostaining in other brain areas of 4.1B KO mice as illustrated in the somato-sensory cortex (Fig. S2). However, a precise quantitative study would be required to examine whether selective dysmyelination of GABAergic interneurons is induced by 4.1B deficiency throughout cortical layers and in particular, in layer2/3, in which nearly half of the myelin ensheaths inhibitory neurons (Micheva et al., 2016; Stedehouder et al., 2017). On the other hand, the band 4.1 transcripts undergo extensive alternative splicing and belongs to a family of proteins including 4.1G, 4.1N, and 4.1R proteins expressed in brain (Buttermore et al., 2011; Ivanovic et al., 2012; Kang et al., 2009). Therefore, we can hypothesize that the cell-type specific effect of 4.1B deletion may rely on compensatory mechanisms such as upregulation of 4.1G or 4.1R, as shown in the PNS (Einheber et al., 2013; Horresh et al., 2010).

### 4.1B is required for the assembly of nodal and juxtaparanodal domains in myelinated inhibitory axons

In addition to the severe reduction of myelin sheath coverage, 4.1B deficiency induces alteration of juxtaparanodal and nodal ion channels. The scaffolding protein 4.1B is linked to the Caspr2/Contactin2 cell adhesion complex at juxtaparanodes (Brivio et al., 2017; Buttermore et al., 2011; Zhang et al., 2013; Zonta et al., 2008). As previously reported in the PNS and CNS (Buttermore et al., 2011; Cifuentes-Diaz et al., 2011; Einheber et al., 2013; Horresh et al., 2010), juxtaparanodal Kv1 clustering is strongly reduced in 4.1B-deficient GABAergic axons of the hippocampus. We observed a similar alteration in *Contactin2*^-/-^ and *Caspr2*^-/-^ GABAergic axons, although some residual expression of Kv1.2 can be detected at the juxtaparanodes of the mutant mice. The Kv1 channels are localized under the compact myelin and their function is still elusive under physiological conditions (Pinatel and Faivre-Sarrailh, 2021). On the other hand, protein 4.1B binds Caspr at paranodes where it organizes the boundary with the nodal Nav channels. Strikingly, the nodal and heminodal Nav channels are no longer juxtaposed to the paranodes in the adult *4.1B*^-/-^ mice, as previously reported in the developing spinal cord and likely as a consequence of the disorganization of the spectrin cytoskeleton (Brivio et al., 2017).

Whether the loss of myelin in the 4.1B mutant influences ion channel expression at the AIS is another intriguing question. We observed structural modification of the AIS of SST cells in the stratum oriens. The length of the AIS as measured with AnkyrinG is significantly shorter in 4.1B KO by comparison with wild-type mice, whereas its onset from the soma or the primary dendrite is not modified. Protein 4.1B is not required for the stabilization of AnkyrinG at the AIS as shown in cultured hippocampal interneurons from *4.1B*^-/-^ mice (Bonetto et al., 2019). However, 4.1B interacting with NuMA1 inhibits endocytosis of AIS membrane proteins and is required for AIS assembly, but not maintenance, as shown by shRNA silencing 4.1B expression at early stage of hippocampal culture (Torii et al., 2020). Therefore, we can propose that the AIS shortening in 4.1B-deficient SST interneurons may not be a primary event but may result from an adaptation to the reduction of myelin axonal coverage. Adaptation of the AIS has been reported in the cuprizone chemical model of demyelination (Hamada and Kole, 2015).

### Class-specific effect of dysmyelination on the excitability of hippocampal interneurons

A defining feature of inhibitory interneuron subsets is their precise axonal arborisation whereby inhibitory synapses target specific subdomains of pyramidal neurons. PV basket cells innervate the perisomatic region and the SST O-LM interneurons connect the distal dendrites of pyramidal neurons. We observed a dramatic loss of myelin in the stratum radiatum whereas myelination appeared slightly affected in the stratum pyramidale. Such a layer-specific alteration may indicate that the O-LM and bistratified interneurons may be more affected than the basket cells. The myelinated pattern of O-LM SST interneurons is highly visible since they extend myelinated axons from the stratum oriens straight across the stratum radiatum. The sheath coverage of O-LM axons is dramatically reduced by 55% in the 4.1B KO by comparison with wild-type mice.

We asked whether dysmyelination may induce changes in the neuronal excitability of interneuron subtypes innervating the different layers of the CA1 hippocampus. In this context, we showed that PV basket cells displayed a similar pattern of fast-spiking discharge in *4.1B*^-/-^ than in control mice. Even if the rheobase tended to decrease suggesting that *4.1B*^-/-^ basket cells could be more excitable than controls, we found no change in the other intrinsic parameters detected at the soma. In contrast, the dysmyelinated phenotype of O-LM interneurons is associated with lower excitability in *4.1B*^-/-^ mice. This was correlated with a significant decrease in the input resistance and a significant increase in the AHP area. Such electrophysiological modifications may reflect changes in ion channel expression at the AIS. In cortical pyramidal neurons, demyelination induced by cuprizone causes restructuring of AIS and reduces excitability at this site. These acutely demyelinated axons displayed Kv7.3 channel expression extended at the distal AIS (Hamada and Kole, 2015). The *I*_M_ current comprised of K_V_7 channels is known to control the interevent intervals and is particularly effective in influencing firing frequency of O-LM cells (Lawrence et al., 2006). Therefore, one can hypothesize that the increase of AHP area in *4.1B*^-/-^ O-LM cells could be due to a change in Kv7 expression at the AIS. In addition, we showed that the length of the AIS is significantly reduced without any change in its distance from the soma, which could lead to a decrease in excitability. Indeed, structural characteristics of the AIS, such as the length and/or distance to the soma, strongly affect the excitability and firing behavior of neurons (Kole and Stuart, 2012; Kuba, 2012; Yamada and Kuba, 2016). Along this line, the derivative analysis suggests that the AP is initiated closer to the soma in *4.1B*^-/-^ O-LM neurons than in controls. In addition, protein 4.1B is associated in a complex with the Kv1 channels along the axon. The clustering of Kv1 channels is altered at juxtaparanodes of 4.1B-deficient mice and as a consequence, Kv1 can be diffuse along the axon to participate in the modulation of intrinsic excitability (Rama et al., 2017).

Functionally, it is well established that PV interneurons are involved in perisomatic inhibition while SST interneurons take part in dendritic inhibition (Somogyi and Klausberger, 2005; Udakis et al., 2020). Thus, we investigated whether the inhibitory inputs could be modified onto the pyramidal cells. We showed that the occurrence probability of small amplitude sIPSCs is reduced in *4.1B*^-/-^ mice suggesting that the distal and not the proximal inhibitory inputs onto pyramidal cells may be decreased as a consequence of the reduced excitability of O-LM cells. The mIPSCs recorded in pyramidal cells did not differ in amplitude and frequency between *4.1B*^+/+^ and *4.1B*^-/-^ neurons suggesting that the number of presynaptic terminals is similar between both genotypes. This last result indicates that dysmyelination of inhibitory axons in CA1 may not be associated with structural synaptic alterations, but a reduction of distal inhibitory inputs.

An important point to keep in view is that myelin sheath, apart from its insulator role, provides metabolic and trophic support in the central nervous system (Kann et al., 2014; Krasnow and Attwell, 2016). As a possible consequence of altered trophic support, it has been reported that in cortical neurons from lysophosphatidylcholine demyelinated mice, there is a decrease in mIPSC amplitude and frequency consistent with a selective loss of PV inhibitory synapses (Zoupi et al., 2021). Similarly, demyelination of cortical PV basket cells from cuprizone treated mice strongly reduces the density of inhibitory presynaptic terminals onto the soma of pyramidal cells (Dubey et al., 2022). In contrast, we did not evidence synaptic alterations associated with dysmyelinated interneurons in 4.1B KO mice. As for long projecting neurons, it is also hypothesized that myelin optimizes the propagation of APs along inhibitory axons. Myelination defects of cortical PV fast-spiking cells are associated with reduced firing frequency and delayed predicted conduction velocity of their APs (Benamer et al., 2020). Indeed, a positive correlation between the degree of axonal myelination and conduction velocity has been reported for cortical PV interneurons (Micheva et al., 2021). It has been also shown that the topography of interneuron myelination depends on the axon diameter and interbranch distance (Stedehouder et al., 2019). Here, we observed that the unramified axonal segment of O-LM cells crossing the stratum radiatum is ensheathed by myelin throughout. The strong reduction of myelin coverage along O-LM cell axons in 4.1B KO mice should decrease AP conduction velocity. In acute demyelinated cortical axons, conduction velocity of APs is reduced and ectopic APs appear; this is associated with a redistribution of Na^+^ channels at branch points with increased nodal expression length of Nav1.6 (Hamada and Kole, 2015). Here, we show mislocalization and increased length of nodal Nav channels that could induce modifications in membrane capacitance and axial resistance to current flow from the node into the internode (Arancibia-Cárcamo et al., 2017). The misdistribution of ion channels implies that AP propagation should be altered along dysmyelinated O-LM axons in *4.1B*^-/-^ mice.

### Selective myelin alteration of hippocampal interneurons is associated with impairment of spatial working memory

Myelination of interneurons may play a critical role in brain functions such as learning and memory (Bonetto et al., 2021; Yang et al., 2020). A dysfunction of hippocampal inhibitory neurons can alter theta and gamma oscillations involved in navigation performance (Kalemaki et al., 2018; Kunz et al., 2019; White et al., 2012). The hippocampal SST cells have been reported to contribute to learning-induced persistent plasticity and spatial memory consolidation (Honoré et al., 2021). For example, the selective loss of mTORC1 function in SST cells impaired spatial long-term memory as tested through the Barnes maze with a decrease in search strategy and performance (Artinian et al., 2019). In the present study, we observed that the spatial working memory is affected in 4.1B KO mice as tested through the Y-maze and Barnes maze tests. In particular, using the Barnes maze, we observed that 4.1B-deficient mice displayed altered performance and spatial search strategy during learning trials and probe test. The O-LM cells are modulating the afferents onto pyramidal cell dendrites, facilitating through disinhibition the Schaffer collateral pathway from CA3 and down-regulating the temporo-ammonic pathway from the entorhinal cortex (Leão et al., 2012). The severe loss of myelin in interneurons of the hippocampus may be associated with a dysfunction in the inhibitory network. A slight apparent disorganization of myelin was also observed in the molecular layer of the dentate gyrus, likely corresponding to inputs from the entorhinal cortex. Although we do not exclude a dysmyelination effect of 4.1B deficiency of extra-hippocampal afferences connecting the hippocampus, our results highlight the role of myelin in O-LM interneurons in fine tuning the hippocampal inhibitory drive that may be involved in spatial exploration and memory.

This study also addresses the contribution of dysmyelination in interneuropathies described in multiple sclerosis. Demyelinated lesions have been reported in the hippocampus of multiple sclerosis patients correlated with cognitive alterations (Geurts et al., 2007). GABAergic interneurons are selectively vulnerable to demyelination in multiple sclerosis (Zoupi et al., 2021). Inhibitory network modifications could lead to an increased incidence of epileptic seizures in multiple sclerosis patients (Kelley and Rodriguez, 2009) and are associated with diminished cognitive function (Nicholas et al., 2016). Here we show a subtype-specific role of myelin in hippocampal GABAergic interneurons. This raises the intriguing question of whether subtypes of interneurons may be specifically affected in demyelinating diseases.

## METHODS

### Animals

The care and use of mice in all experiments were carried out according to the European and Institutional guidelines for the care and use of laboratory animals and approved by the local authority (laboratory’s agreement number D13-055-8, Préfecture des Bouches du Rhône). The following mouse strains were used in this study: C57bl/6 mice (Janvier Breeding Centre), previously described *Tag-1*/*Contactin2* KO mice (Traka et al., 2003), *4.1B* KO mice (Cifuentes-Diaz et al., 2011) and *Cntnap2*/*Caspr2* KO mice (Poliak et al., 2003). Both *Contactin2* and *4.1B* KO mice were crossed with *Lhx6-Cre* (strain 026555) and *Ai14* (strain 007908) mouse strains (Jackson Laboratory, Bar Harbor, ME, USA) to obtain mice with the following genotypes *Contactin2*^-/-^;*Lhx6-Cre;Ai14* and *4.1B*^-/-^;*Lhx6-Cre;Ai14* expressing tdTomato. Fixed brains from *Caspr2* KO obtained from the Jackson laboratory (strain 017482) were a gift from Dr. N. Noraz (NeuroMyogene Institute, Lyon, France).

### Immunofluorescence staining, confocal microscopy and image analysis

Rabbit antiserum against protein 4.1B (Cifuentes-Diaz et al., 2011) and rabbit anti-Caspr (Bonnon et al., 2007) were described previously. The following commercial primary antibodies were used: rabbit anti-Olig2 antibody and mouse anti-panNav mAb (clone K58/35) purchased from Sigma-Aldrich Chimie, (Saint-Quentin-Fallavier, France), rat anti-MBP mAb (ab7349) from Abcam (Cambridge, UK), goat anti-PV antibody (PVG-214) from Swant (Burgdorf, Switzerland), goat anti-SST mAb (sc55565) and mouse anti-SST (sc55565) from Santa-Cruz Biotechnology (Heidelberg, Germany). Mouse anti-ankyrinG (clone N106/36), anti-Kv1.2 (clone K14/16), anti-Kv3.1b (clone N16B/8) mAbs were obtained from NeuroMab (UC Davis/NIH NeuroMab Facility). AlexaFluor-405, 488, −568 and −647-conjugated secondary antibodies were purchased from Molecular Probes (Life Technologies, Courtaboeuf, France).

P25, P30, P35, and P70 mice were deeply anesthetized with a mix of Zoletyl/Domitor and then transcardially perfused with PBS followed by 2% paraformaldehyde in PBS. Brains were removed and placed in the same fixative overnight. Eighty micron-thick vibratome sections were permeabilized and blocked for 1 h in PBS containing 5% horse serum and 0.3% Triton X-100. Antigen retrieval in 10 mM citrate buffer, pH6, at 90°C for 30 min was performed before immunostaining with anti-panNav and anti-AnkyrinG antibodies. Floating sections were incubated for 2 days at 4°C with combinations of the following primary antibodies: goat anti-PV (1:1000), mouse anti-SST (1:200), goat anti-SST (1:500), rabbit anti-Caspr (1:2000), mouse anti-Kv1.2 (1:400), mouse anti-Kv3.1b (1:100), mouse anti-ankyrinG (1:100), mouse anti-panNav (1:200), rat anti-MBP (1:200), rabbit anti-Olig2 (1:2500). Sections were then washed and incubated with the appropriate AlexaFluor-conjugated secondary antibodies (1:500) overnight, washed and mounted on slides with Vectashield mounting medium (Vector Laboratory, Eurobio Scientific, Les Ulis, France). Image acquisition was performed on a Zeiss (Carl-Zeiss, Iena, Germany) laser-scanning microscope LSM780 equipped with 40X 1.32 NA oil-immersion objective or 20X objective. Images of AlexaFluor-stained cells were obtained using the 488 nm band of an Argon laser and the 405 nm, 568 nm and 647 nm bands of a solid-state laser for excitation. Fluorescence images were collected automatically with an average of two-frame scans at airy 1. Maximum intensity projection of images was carried out using ImageJ software (NIH). Images are single confocal sections unless the number of z-steps is indicated.

The density of paranodes immunolabeled for Caspr was estimated in the different layers (stratum oriens, stratum pyramidale, stratum radiatum and stratum lacunosum-moleculare) of the CA1 hippocampus at P70. The density of single paranodes (1 doublet of paranodes=2 single paranodes) was quantified in control and 4.1B-deficient mice using maximum intensity projection images (5-z steps of 2 μm) (ROI of 10^5^ μm^2^/animal, n=3-4 mice/genotype). The total length of myelinated axons was measured in the stratum pyramidale (1 bin) and stratum radiatum divided in 5 bins (40×300 μm) on single confocal sections, n=3-4 mice/genotype. We also measured the total length of Lhx6- or PV-positive myelinated axon in the stratum radiatum at P35 or P70 (5 ROIs of 10^5^ μm^2^ from 3 mice/genotype). The length of myelinated SST axons was measured in the stratum radiatum of control and 4.1B KO mice at P35 (stacks of 1 μm 5-8z steps; 10 ROIs of 10^5^ μm^2^ from 3 mice/genotype). To analyze the extent of myelin coverage of individual SST-positive axons across the stratum radiatum, only axons with a length>100μm were taken into account (n=29-35 axons from 3 mice/genotype). Three-D reconstruction of myelinated axons was performed using the Neurolucida Explorer Software (Micro Bright Field, Inc., Williston, VT, USA). The distribution of the lengths of internodes was evaluated in the CA1 hippocampus of control and *4.1B*^-/-^ *Lhx6-Cre;tdTomato* mice at P70. Double-immunostaining for MBP and Caspr and confocal imaging were performed with analysis on maximum intensity projection images (25 to 50 μm-width z-stacks). We measured the length of myelin sheaths bordered at each end by a paranode in 4 ROIs of 10^5^ μm^2^ from 2 mice/genotype (n= 275 in control and n=191 in *4.1B*^-/-^ mice). The length of nodal Nav was measured in the CA1 hippocampus of control *Lhx6-Cre;tdTomato* at P30 and P70. Triple-immunostaining for Caspr, MBP and panNav was performed and Lhx6-positive axons expressing tdTomato were detected with the red channel. The distribution of nodal length was analyzed on confocal images (1 μm 2-5z-steps) from 2 mice/age (P30: n=41, P70: n=64). Next the length of nodal or heminodal Nav channel clusters and the gap between Nav clusters and paranode were measured in the CA1 hippocampus of *4.1B*^+/+^ and *4.1B*^-/-^ *Lhx6-Cre;tdTomato* mice at P70 (n>25/genotype from 2-3 mice/genotype). The clustering of juxtaparanodal Kv1 channels was estimated in the CA1 hippocampus of wild-type, *4.1B*^-/-^, *Caspr2*^-/-^, and *Contactin2*^-/-^ mice at P70. The percentage of single paranodes bordered by Kv1.2 immunostaining in PV-positive axons was quantified on 5-10 μm z-stack confocal images. More than 150 single paranodes were analyzed in each genotype in 7-11 ROIs from 3 mice/genotype). Mutant mice were compared with their respective controls in double-blind experiments.

### Electrophysiological recordings

Mice were deeply anesthetized with Zoletyl/Domitor and subsequently decapitated. Brains were quickly removed from the skull and placed into ice-cold oxygenated (95% O2 and 5% CO2) cutting solution containing the following (in mM): 132 choline, 2.5 KCl, 1.25 NaH2PO4, 25 NaHCO3, 7 MgCl2, 0.5 CaCl2, and 8 D-glucose (pH: 7.4, 290–300 mOsm/L). 300 μm-thick coronal brain slices were prepared using a VT1200S microtome (Leica Microsystems, Wetzlar, Germany). Slices were left at room temperature in oxygenated (95% O2 and 5% CO2) solution of artificial cerebrospinal fluid (aCSF) containing the following (in mM): 126 NaCl, 3.5 KCl, 1.2 NaH2PO4, 26 NaHCO3, 1.3 MgCl2, 2.0 CaCl2, and 10 D-glucose, pH 7.4. Then, slices were transferred into a submerged recording chamber and perfused with oxygenated aCSF at a flow rate of 2 to 3 ml/min. Recordings were performed under visual guidance using infrared differential interference contrast microscopy (SliceScope Pro 3000M, Scientifica, Uckfield, UK). tdTomato in neurons was excited by a UV lamp and the fluorescence was visualized using a CCD camera (Hamamatsu, Japan). Whole-cell patch-clamp recordings were made using a Multiclamp 700B amplifier (Molecular Devices, CA, USA), filtered at 2 kHz using the built-in Bessel filter of the amplifier. Data were digitized at 20 kHz with a Digidata 1440A (Molecular Devices) to a personal computer, and acquired using Clampex 10.1 software (PClamp, Molecular Devices). For current clamp recording mode patch pipettes were pulled from borosilicate glass tubing with resistances of 6–8 MΩ and filled with an internal solution containing the following (in mM): 130 KMeSO4, 5 KCl, 10 4-(2-hydroxyethyl)-1-piperazi-methanesulfonic acid, 2.5 MgATP, 0.3 NaGTP, 0.2 ethyleneglycoltetraacetic acid, 10 phosphocreatine. Access resistance ranged between 15 and 50 MΩ, and the results were discarded if the access resistance changed by >20%. Neurons recorded in hippocampal CA1 region were held at a holding potential of −70 mV. Series resistance (between 5–21 MΩ) was monitored throughout each experiment and neurons with a change in series resistance of more than 25% were excluded from the analysis.

To study intrinsic electrophysiological properties of CA1 inhibitory neurons, we analyzed the voltage responses to a series of hyperpolarizing and depolarizing square current pulses of 500 ms duration of amplitudes between −70 pA and 130 pA with 10 pA step intervals from a holding potential of −70 mV in each cell. The voltage responses of neurons upon the current injections also helped to distinguish interneurons of a fast-spiking character having no or small ‘sag’ response to a hyperpolarizing current pulse from cells with lower spiking rate and a larger ‘sag’, typical of O-LM cells (Maccaferri and McBain, 1996) (Table 1 and 2). Input resistance (Rinput) was determined by calculating the slope of the plot of the membrane potential variation induced by a hyperpolarizing 500 ms step of current (I: from −70 pA to 0 pA). Firing frequency was studied by injecting 500 ms pulses of depolarizing current (I: from 10 pA up to 130 pA) into the cell and plotting the spike frequency (f) as a function of the current intensity (f/I plot). For presumed fast-spiking neurons, a second protocol was used were current steps varied from +100 pA to +400 pA with 100 pA steps, in order to analyze possible differences in high-speed discharge. Both analyses were done using Prism 8 software (GraphPad Software, La Jolla, CA). For the analysis of AP, the first AP evoked by a suprathreshold depolarizing current pulse (considered as the Rheobase) was selected. The following AP features were calculated using Minianalysis software: peak amplitude, area, half-width, threshold, AHP area and AHP peak (Table 1 and 2). We further analyzed the shape of the AP rising phase using multiple derivatives of the raw membrane potential signal using Clampfit software (PClamp, Molecular Devices). In some dV/dt curves appeared a shouldering. A “double bump” in the d^2^V/dt^2^ curve and a value equal to or less than zero of the d^3^V/dt^3^ trace between the two bumps indicated that the shouldering corresponded to a stationary inflection in the rising slope of the AP. This indicated that somatic and axonal components of the AP could be temporally differentiated (Kress et al., 2008; Paterno et al., 2021).

GABA-induced spontaneous inhibitory post synaptic currents (sIPSCs) were recorded in pyramidal cells in voltage-clamp mode using patch pipettes of 8-10 MΩ filled with an internal solution containing the following (in mM): 140 CsCl, 1 MgCl2, 10 HEPES, 4 NaCl, 2 Mg-ATP, 0.3 Na-GTP, 0.1 EGTA. The GABA-mediated current was pharmacologically isolated in the presence of AMPA and NMDA receptor antagonists (10 μM CNQX, 40 μM D-APV respectively). For recording of miniature IPSCs (mIPSCs) TTX (1μM) was added to the ACSF. CNQX, D-APV and TTX were purchased from Tocris Bioscience (Bristol, UK).

### Behavioral analysis

We examined spatial learning memory through Y-maze and Barnes maze from 3-month mice. Heterozygous mice (*4.1B*^+/-^; *Lhx6-Cre; Ai14*) were crossed to generate the following genotypes *4.1B*^+/+^; *Lhx6-Cre; Ai14* and *4.1B*^-/-^; *Lhx6-Cre; Ai14.* Animals were first handled 2 minutes per day for 4 days. *Spontaneous Y-maze test:* Animals (4 males and 4 females of each genotype) were placed in the center of a Y-maze (Noldus apparatus, Wageningen, The Netherlands) and no external cues were visible from inside the maze. Spontaneous alternation was recorded during 8 min (d’Isa et al., 2021; Deacon and Rawlins, 2006). *Barnes maze test:* Animals of each genotype were tested in the maze in random order (controls: 11 males and 3 females; 4.1B KO: 7 males and 3 females). Visual cues were placed around the maze, and the ceiling was equipped with high illumination (120 lux) and background sound (85 dB). The training for the test was performed for 4 days. Each animal adapted to an escape box for 2 min on the first day, was placed in the center of the maze and explored the maze to find an escape hole, which is connected to the escape box. Once the animal entered the escape box, it was left there for 30 s, and the trial ended. Each animal was given 2 daily trials with a 30 min inter-trial interval for the 4 consecutive days. The animals who did not find the escape box after 120 s were gently guided toward the escape hole and left there for 30 s. The retention test was performed at 1 day after the training. The escape box was removed, and the latency time to reach the entry zone, where the escape box was previously located, was recorded during 120 s for each animal. Using an automatic tracking system, the trajectory was recorded (Ethovision, Noldus). The path length, average speed, latency to reach or to enter the escape hole were quantified. Decreases in latency to escape hole with increased training is indicating that spatial learning occurred. We estimated the percentage of trials using spatial (direct, corrected, with long-correction) or non-spatial (random or serial) search strategies (Cheng et al., 2019; Illouz et al., 2016).

### Statistical analysis

All values are given as means ± SEM. All statistical tests were performed using Prism 8 (GraphPad Software). For comparisons of two independent groups, we used Mann–Whitney test or Student’s t-test when values lie in a normal distribution using Shapiro-Wilk test. For multiple group comparisons, we used two-way ANOVA followed by the Sidak’s multiple comparison test. To quantify the probability that two sets of samples were drawn from the same probability distribution, we used the Kolmogorov-Smirnov test. To compare the rate of SST cells with stationary inflection to those without, we used Fisher’s exact test.

## Acknowledgments

We are grateful to Dr Thomas Marissal for helpful discussions. We thank the University of California Davis/National Institutes of Health NeuroMab Facility and Developmental Studies Hybridoma Bank of the University of Iowa. This work was supported by the Association pour la Recherche sur la Sclérose en Plaques (ARSEP) to DP, CFS and DK.

