## Supplemental Figures 1-4 for "A class-specific effect of dysmyelination on the excitability of hippocampal interneurons"

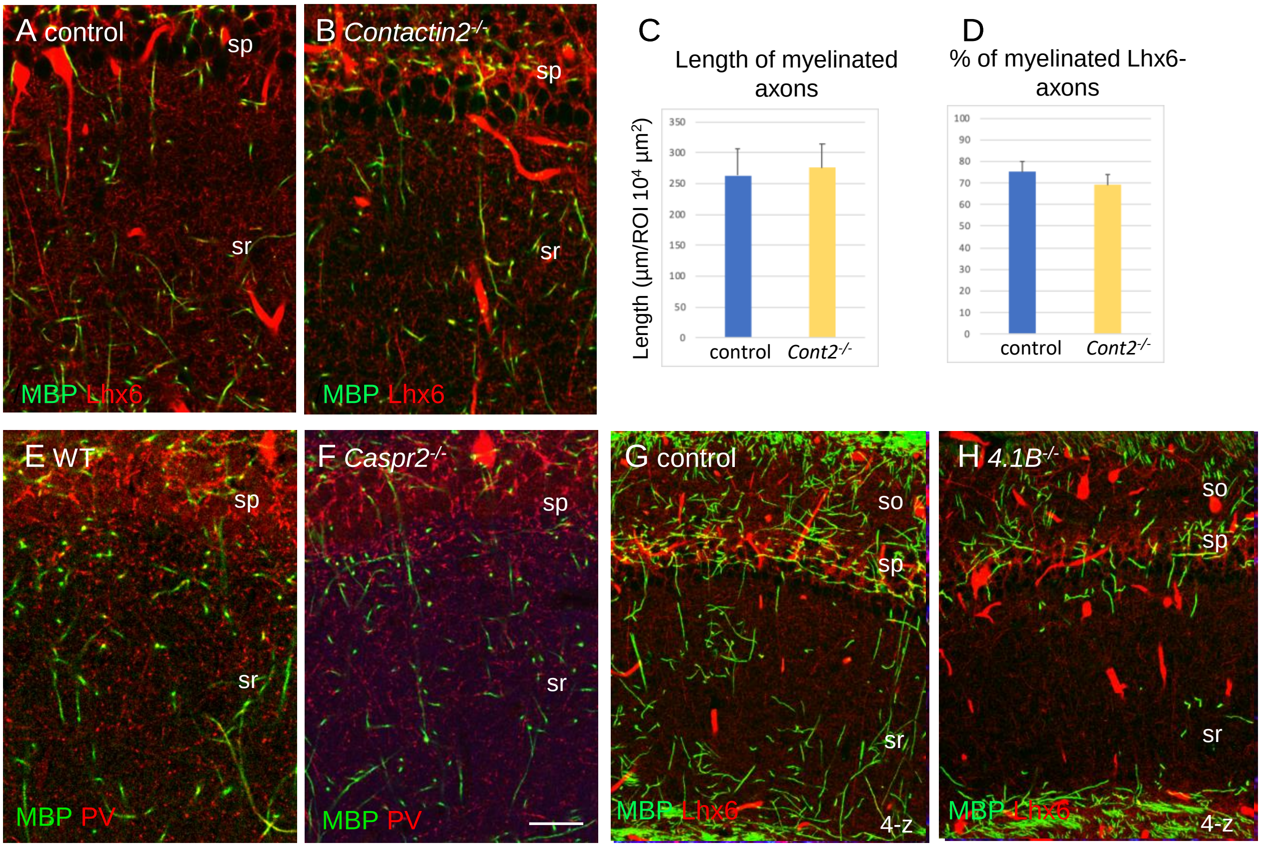


**Figure S1: Myelin pattern in the CA1 hippocampus of *Contactin2*$^{-/-}$, *Caspr2*$^{-/-}$, and *4.1B*$^{-/-}$ mice**

CA1 hippocampus stained for MBP (green). A, B, G, H: *Lhx6-Cre;tdTomato* mice: control (A, G), *Contactin2*$^{-/-}$ (B) and *4.1B*$^{-/-}$ (H) mice at P70 showing axons of PV and SST cells expressing the tdTomato (red). Note that the blood vessels are fluorescent. Single confocal image (A, B) or maximum intensity of confocal images of 4-z steps of 2 µm (G, H). E, F: CA1 hippocampus of wild-type and *Caspr2*$^{-/-}$ mice at P70 immunostained for MBP (green) and PV (red). Bar: 20 µm. C, D: Quantification of total myelin length/$10^4$µm$^2$ in the stratum radiatum of control and *Contactin2*$^{-/-}$ mice and percentage of myelinated Lhx6-expressing axons at P35. Mean ± SEM of 3 mice/genotype. No significant difference by comparison with control (P=0.4 and P=0.3, respectively; Mann-Whitney test).

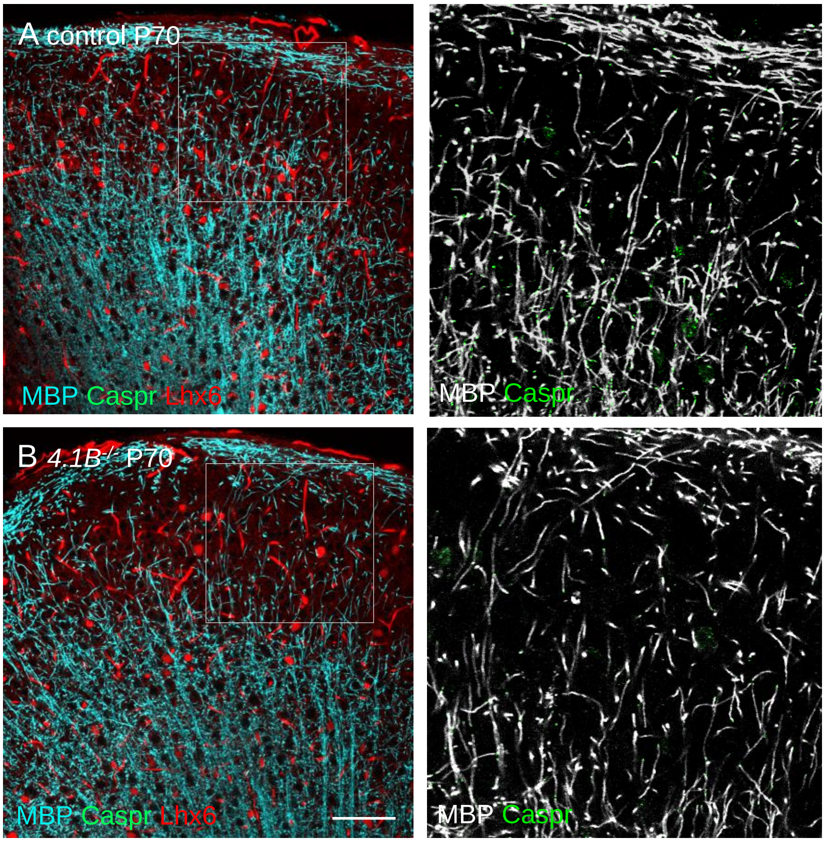


**Figure S2: Myelin pattern is preserved in the cortex of *4.1B$^{-/-}$* mice at P70**
Coronal brain sections showing the somato-sensory cortex of *Lhx6-Cre;tdTomato* control (A) and *4.1B$^{-/-}$* (B) mice at P70. Double-staining for MBP (Cyan) and Caspr (green). Note in the inset that the myelin pattern (grey) is preserved in the cortical layer 2/3 of *4.1B$^{-/-}$* mice. Bar: 100 µm.

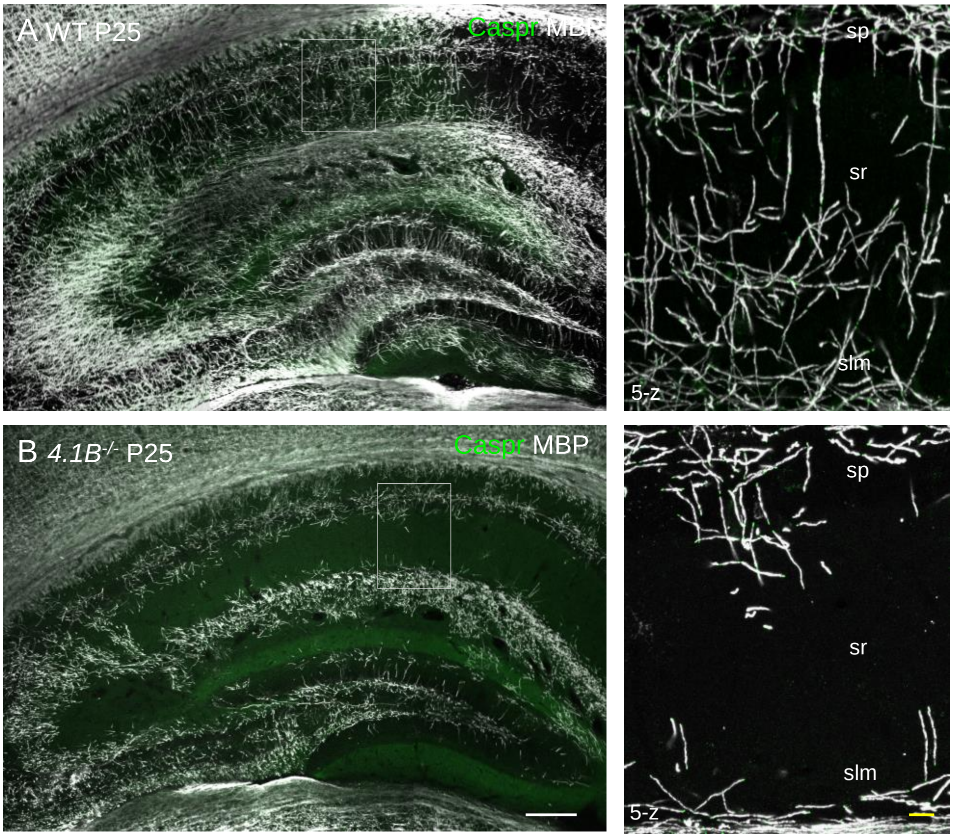


**Figure S3: Myelin pattern alteration in the hippocampus of *4.1B$^{-/-}$* mice at P25**
Hippocampus from P25 wild-type and *4.1B$^{-/-}$* mice immunostained for MBP (grey) and caspr (green). Note the dramatic loss of myelin sheaths in the CA1 stratum radiatum of the *4.1B$^{-/-}$* hippocampus. Bar: 200 µm in A, B; 25 µm in the insets.

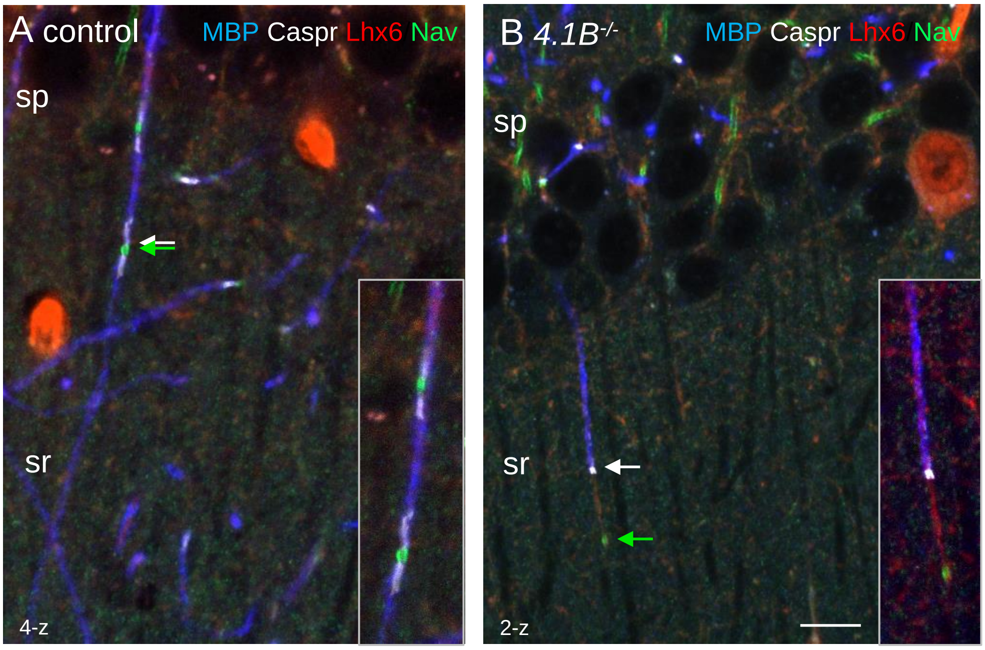


**Figure S4: Myelination and positioning of Nav channels is altered in 4.1B-deficient GABAergic axons**

Hippocampal sections of P70 *Lhx6-Cre;tdTomato* control and *4.1B*$^{-/-}$ mice were immunostained for MBP (blue), Caspr (white) and panNav (green). tdTomato expressed in Lhx6-positive cells is in red. A: a control myelinated Lhx6-positive axon crossing the stratum radiatum showing nodes of Ranvier (A, arrows). B: a partially myelinated Lhx6-positive axon displays a large gap between the Nav cluster and heminode in 4.1B-deficient hippocampus (B, arrows). Confocal images with maximum intensity of 2-z or 4-z steps of 1 µm. Bar: 10 µm.
